# Fine-tuning neuron number: *Lzts* as a key regulator of neurogenesis and cell proliferation in *Ciona*

**DOI:** 10.64898/2026.09.24.754029

**Authors:** Paola Olivo, Ugo Coppola, Mark Passero, Christopher J. Johnson, Sabrina A. Hernandez, Alberto Stolfi, Filomena Ristoratore

**Author notes:** these authors equally contributed to the paper (Paola Olivo, Ugo Coppola).

## Abstract

How progenitor mitotic timing is coupled to the number of neurons in a nervous system remains poorly understood, in part because most model systems have large, variable progenitor pools that obscure this relationship. The non-vertebrate chordate *Ciona robusta* offers a tractable alternative: its larval nervous system arises from an invariant lineage, including four Bipolar Tail Neurons (BTNs) that are reproducibly generated from the caudal neural plate border. Here we investigate the developmental function of the single *Ciona* ortholog belonging to the leucine zipper tumor suppressor (*Lzts*) gene family, whose vertebrate members are implicated in cell-cycle regulation and tumor suppression. In *Ciona*, *Lzts* is expressed in several embryonic tissues, including differentiating BTNs. Tissue-specific CRISPR/Cas9 knockout of *Lzts* in the BTN lineage produced supernumerary BTNs, without expanding the embryonic Neurog+ domain, indicating that Lzts acts downstream of BTN specification to limit neuron number. Lzts1 promotes Cdk1 activity via stabilization of Cdc25C, co - expression of *Ciona* Cdc25C or a non-phosphorylatable Cdk1/2/3 mutant rescued normal BTN number in *Lzts* crispants, suggesting the existence of an ancestral Lzts-Cdc25-Cdk1 circuit controlling neural cell cycle. These findings establish *Ciona* Lzts as a regulator of mitotic tempo determining the BTN number and suggest that this cell-cycle regulatory axis links proliferation to neurogenesis across chordates *in vivo*.

## Introduction

Neurogenesis is a sequence of cellular and physiological modifications in which different pathways interact with each other. However, the majority of these developmental pathways and their molecular mechanisms are still poorly understood. A central question is how the duration and number of progenitor divisions are coupled to the final number of neurons produced. In systems with large, heterogeneous progenitor pools, such as the mammalian neocortex, this coupling is obscured by long developmental times and considerable variability across individuals, making it difficult to isolate the direct contribution of mitotic kinetics to final cell number. The invertebrate chordate *Ciona robusta* represents a useful and tractable model to study the regulation of cell behaviour during neurogenesis^1,2^. Its larval nervous system originates from a small, essentially invariant lineage tree, in which each precursor division and its progeny can be tracked individually. From the caudal part of the lateral borders of the neural plate, a type of sensory relay neurons, the Bipolar Tail Neurons (BTNs) are derived^1^. Exactly four BTNs are invariantly located along the tail nerve cord, two on either left/right side, each characterized by two long processes that extend in opposite directions along the anterior-posterior axis. BTNs are further sub-classified as GABAergic anterior BTNs (aBTNs) and cholinergic posterior BTNs (pBTNs), derived from adjacent lineages^1,3^. This makes the BTN lineage an unusually direct readout for any manipulation that alters the timing or outcome of proliferation: extra or missing divisions translate into readily obvious extra or missing neurons.

Several features of BTNs neurons such as their developmental origin, expression of different transcription factors and early *Neurogenin* (*Neurog)* and *Hmx* expression, together with their proposed role in relaying peripheral sensory inputs to the central nervous system (CNS), suggest that the BTNs are homologs of the vertebrate cranial sensory neurons^4^. Previous studies on BTN specification showed that *Neurog* expression is necessary and sufficient for the specification of BTNs from the caudal neural plate borders^1,2^, controlling the activation of multiple downstream genes. Among these, the leucine zipper tumor suppressor (*Lzts*) gene was identified as upregulated by Neurogenin^2^. Members of the leucine zipper tumor suppressor (Lzts) protein family are thought to function in cell cycle control^5^. In cancer cells, Lzts1 and Lzts2 inhibit cell growth and suppress tumorigenesis, and *Lzts1* mutations are involved in several human neoplasias^5,6^. In mammals, *Lzts1,* also known as *FEZ1*^7^, is upregulated by Neurogenin family factors and acts as a master modulator of neurogenic cell delamination^8^. Furthermore, both mouse and chick Lzts1 orthologs are expressed in the spinal cord during development^9^. Additionally, the zebrafish *lzts2* gene regulates cell movements associated with Wnt signaling^10^. In mammals, *Lzts2* mutants show clear urinary and kidney defects during embryonic development^11^. Intriguingly, recent analyses demonstrated a central role for *Lzts3* (aka *Prosapip*) in the context of synaptic development^12^. Furthermore, mammalian *Lzts4* (aka *Nb4bp3*) is implicated in multiple types of carcinoma^13,14^ and regulates neuronal and dendritic growth in amphibians^15^.

Given the established roles of vertebrate *Lzts* genes in tumor suppression, and neural development, the analysis of *Ciona Lzts* during BTN formation offers a unique opportunity to investigate how neurogenic transcriptional programs are coupled to the control of neuronal production. As *Ciona* retains a single ancestral *Lzts* ortholog within an invariant neurogenic lineage, it provides a particularly tractable system to explore the ancestral functions of the Lzts family in chordate nervous system development.

Here, we reconstruct the evolutionary history of the *Lzts* gene family across metazoans and investigate the function of the single *Lzts* ortholog present in *Ciona*. Combining tissue-specific CRISPR/Cas9-mediated knockout with specific markers, we show that *Lzts* controls the precise number of BTNs generated during development and acts downstream of *Neurog*. Loss of *Lzts* resulted in supernumerary BTNs, indicating a role in restricting neuronal production rather than BTN specification itself, through a conserved interaction with Cdc25-Cdk1 cell cycle regulators^5,16^. Overall, our data pave the way to connecting a conserved Neurog-dependent gene regulatory network (GRN) to regulation of cell cycle and delamination, identifying a new player in the development of the BTNs. Here, our work in *Ciona* indicates the Lzts1-Cdc25-Cdk1 axis as a main regulator *in vivo* of mitosis in the chordate nervous system.

## Results

### Phylogenetic reconstruction of *Lzts* family in metazoans

To garner a better understanding of the evolutionary history of the *Lzts* family, we carried out a comprehensive evolutionary analysis, combining diverse comparative genomics approaches. Using 180 family protein sequences, we built a maximum-likelihood (ML) phylogenetic tree with representatives from the entire animal kingdom (**Figure 1, Supplementary Figure 1**). Using the InterProdatabase available on Ensembl, we confirmed that all the retrieved proteins correspond to the Interpro code for Lzts proteins (data not shown). We did not identify Lzts members from early-branching metazoans (placozoans, sponges, ctenophores, cnidarians), which hints at the potential emergence of *Lzts* genes specifically in bilaterians. Our phylogenetic survey clearly highlights the presence of 5 Lzts clades which we named Lzts1/2/3/4, Lzts1, Lzts2, Lzts3, and Lzts4. Importantly, *Lzts1/2/3/4* is present only in invertebrates and lampreys, while gnathostomes (jawed vertebrates) possess *Lzts1*, *Lzts2*, *Lzts3* and *Lzts4* **(Figure 1**). As expected, Lzts from all the surveyed tunicates formed a sister group to vertebrate Lzts sequences, with the exception of the sequence from the highly divergent pelagic and neotenic tunicate *Oikopleura dioica*^17^. Intriguingly, conservation of intron/exon structure (**Figure 1B, Supplementary Figure 2**) and syntenic analysis of *Lzts* loci in multiple species (**Figure 1C**) suggest that an ancestral *Lzts1/2/3/4* gene (which we henceforth call simply “*Lzts”*) gave rise to four extant ohnologs in vertebrates (*Lzts1, Lzts2, Lzts3,* and *Lzts4*), through two events of whole-genome duplication^18–21^. The presence of a single *Lzts* gene in agnathans (jawless vertebrates) is not entirely inconsistent with this scenario, as new evidence suggests that only one round of whole genome duplication occurred at the base of vertebrates^22^. The lack of a second *Lzts* paralog in the extant agnathans surveyed here (*Petromyzon marinus*, *Eptatretus burgeri*) suggests that it may have been lost after the split from gnathostomes. Additionally, the synteny analysis in multiple vertebrates clarified the orthology of the four vertebrate extant subfamilies (*Lzts1*, *Lzts2*, *Lzts3*, *Lzts4*) (**Supplementary Figures 3-6**), suggesting that both the teleost-specific genome duplication (TSGD, 3R)^23^ and salmon-specific genome duplication (SSGD, 4R)^24^ contributed to the expansion of *Lzts* family in teleosts. As expected^25,26^, we registered multiple gene losses in this clade that re-shaped the *Lzts* repertoire among teleost species. In sum, our analysis highlights how the *Lzts* family has been shaped by multiple genomic events during metazoan evolution, with a clear expansion in jawed vertebrates.

**Figure 1.**
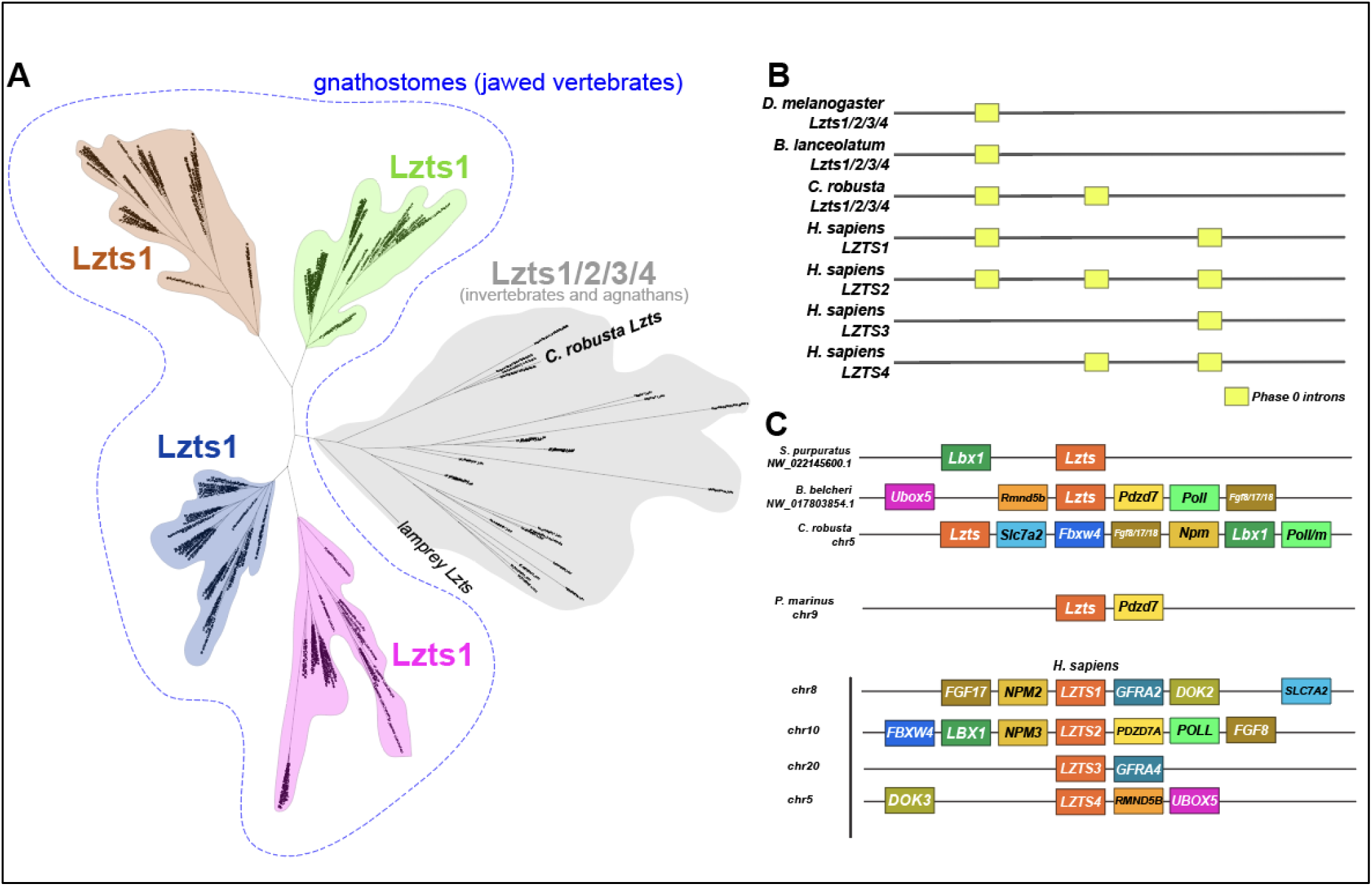
*Lzts* evolution in metazoans. **A)** Evolutionary reconstruction using ML approach, highlighting the presence of five subfamilies, indicated by different colors. Numbers at the branches indicate replicates generated using aLRT method. **B**) Intron code of selected *Lzts* genes in metazoans. Yellow rectangles indicate conserved intron/exon junctions. **C**) Synteny analysis of key invertebrate and vertebrate *Lzts*. Schematization of conserved genomic environments of *Lzts* (red rectangles) in selected species with relative chromosomes/scaffolds information. Flanking orthologous genes are represented employing rectangles of the same color.

### *Lzts1* is expressed in multiple tissues, including the BTNs of *Ciona robusta*

In vertebrate embryos, the expression of *Lzts* genes has been poorly characterized, with few data in zebrafish early development^10^ and in distinct territories of mammals^5,6,11^. The central role of distinct *Lzts* genes in the nervous system development of mammals^8,12^, together with the scarcity of information on invertebrate *Lzts*, prompted us to investigate *Lzts* expression during the development of the tunicate *Ciona robusta* (**Figure 2**), for its key evolutionary position as the sister group to vertebrates within the chordates^27^. Our whole-mount *in situ* mRNA hybridization analysis showed that *Lzts,* the sole *Lzts1/2/3/4* orthologue in the *Ciona* genome, is expressed in epidermal cells in the anterior and posterior tips at late neurula stage until mid tailbud I stage. As development proceeds, from middle tailbud to late tailbud we find expression in the head, at the boundary between the neural plate and papilla, the notochord and the tail tip (**Figure 2A-F**). Expression in the BTNs was already hinted at by RNAseq^2^, and by *in situ* hybridization we also observed modest expression in the migrating BTNs at the mid-tailbud stage (**Figure 2G,H**). Taken together, these data suggest that *Lzts* is dynamically expressed in diverse tissues during *Ciona* embryogenesis, hinting at diversified developmental roles.

**Figure 2.**
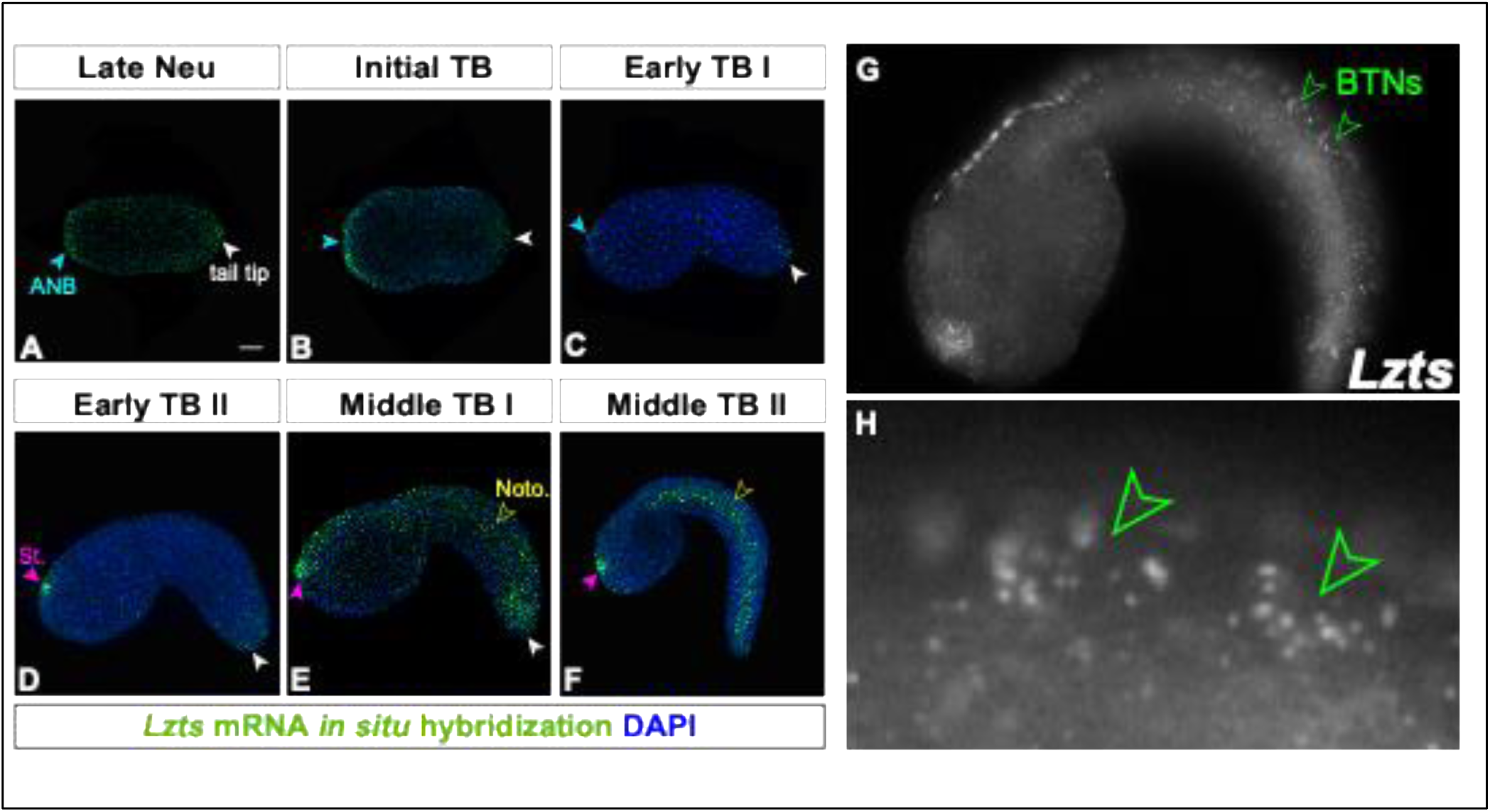
Expression pattern of *Lzts* in *Ciona robusta*. **A-F**) Fluorescent *In situ* hybridization showing dynamic *Lzts* expression from late neurula (A), to Middle tailbud II (F). DAPI has been used as a marker for the non-GFP cells. Expression in epidermal cells of the anterior and posterior tips (from late neurula, blue and white arrows). Expression in notochord, neural papilla and tail tip (yellow arrow). **G, H**) Details of the *Lzts* mRNA localization in BTNs (green arrows). Panels A-F used *in situ* probe from template 1, panels G and H used probe from template 2 (see supplementary file 3 for details).

### *Lzts cis*-regulatory elements expressed in BTNs

In light of the dynamic expression pattern of *Lzts* in *Ciona* embryos (**Figure 2**), we next focused our attention on the putative *cis-*regulatory regions underlying its expression (**Figure 3A**). Hence, we cloned into a GFP reporter vector a region that is highly conserved between *C. robusta* and *C. savignyi* (from −1789 to +6 relative to the start codon). As previously reported for other *cis*-regulatory elements^28,29^, the examined region did not exhibit conservation with vertebrates or other groups of tunicates. We named this reporter plasmid *LztsA*>GFP and electroporated it in *Ciona* zygotes to evaluate its capability to drive GFP expression where endogenous *Lzts* was shown to be expressed. Our results show that *LztsA>GFP* was able to drive expression in the BTNs, in addition to the oral region (stomodeum), papillae, epidermis, notochord, and CNS neurons, all territories of endogenous *Lzts* expression (**Figure 3B, Supplementary Figure 7**). Expression in the oral region appeared to be in both oral ectoderm and underlying oral endoderm (**Figure 3B**). In order to find a *cis-*regulatory element driving specific expression in BTNs cells, we subdivided the whole region into smaller fragments (**Figure 3C-F**). We found that BTN activity peaked around −850 and −500 bp 5’ to the start codon of *Lzts*, though we were unable to fully separate this from *cis*-regulatory activity in other tissues, such as notochord, epidermis, or other neural cells. This suggests the dynamic regulation of *Lzts* in multiple tissues may depend on a complex combination of shared and tissue-specific upstream regulators.

**Figure 3.**
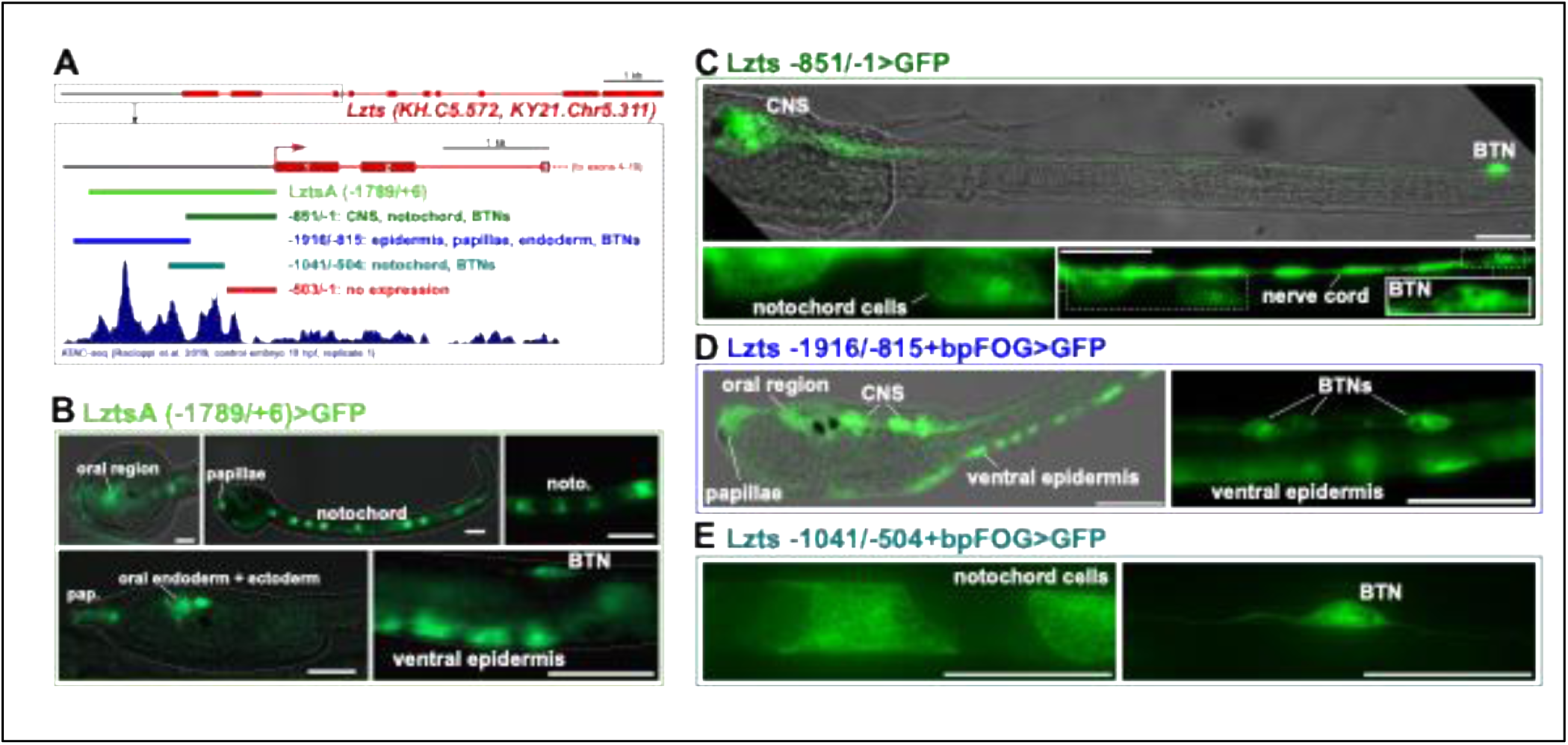
*Cis*-regulatory analysis of *Ciona robusta Lzts*. **A)** Identification of putative *cis*-regulatory elements 5’ upstream of the *Lzts* gene, with a summary of the activities of the genomic fragments used for the regulatory analysis (corresponding to previously published ATAC-seq peaks). **B)** Larvae showing reporter gene expression driven by *LztsA (−1789/+6)>GFP* in oral region (stomodeum/oral ectoderm + oral endoderm), papillae (pap.), notochord (noto.), ventral epidermis, bipolar tail neuron (BTNs). **C)** *Lzts −851/-1>GFP* expression in central nervous system (CNS), notochord, nerve cord and BTNs. **D)** *Lzts −1916/-815 + bpFOG>GFP* driving specific expression in diverse tissues and cell types including BTNs. **E)** *Lzts −1041/-504 + bpFOG>GFP* = basal promoter of *FOG* gene, see methods and Supplementary File 3 for details. Scale bars: 50 μm. All *GFP* sequences except for *LztsA>GFP* tagged with *Unc-76 (Unc-76::GFP)* to label axons efficiently.

### *Ciona Lzts* gene controls normal number of BTNs during development

We were intrigued by the expression of *Lzts* in the BTNs (**Figure 2H**, **Figure 3B-E**), hypothesizing that it may be involved in regulating neurogenesis downstream of Neurog^2^, as Lzts1 does in the mammalian cortex downstream of Neurog1/2^8^. Previously published RNAseq on isolated BTN progenitor cells^2^, revealed a subset of genes upregulated by Neurog, including *Lzts*. In order to clarify the function of *Lzts* in the context of BTN development, we employed a tissue-specific CRISPR/Cas9-based gene editing approach. We designed and selected two single-chain guide RNAs (*sgRNAs)* named *Lzts.384* and *Lzts.441*, with 24% and 25.1% mutagenesis efficiency, respectively (**Supplementary Figure 8**). To perform a phenotypic analysis of *Lzts* crispants in F0 larvae, we used the *Friend of GATA (FOG)* promoter to restrict Cas9 expression to the animal pole which originates the peripheral nervous system, including the BTNs^30,31^. For the electroporations, we combined this with the two selected *sgRNA* expression vectors and a *Rimbp* reporter plasmid (*Rimbp[Intr7A]>GFP)* which strongly labels the BTNs^32^. As a negative control, we employed a non-specific sgRNA expression vector (*U6>Control),* designed not to cut any *Ciona* genome sequence^31^. Strikingly, we observed a significant increase in BTN number in the knockdown condition, with some larvae displaying more than 4 GFP+ BTNs (**Figure 4A,B**). Although each larva invariantly has 4 BTNs, due to reporter plasmid mosaicism, normally fewer than 4 BTNs are labeled by GFP reporters in any given larva. Thus we interpreted the shift to more GFP+ BTNs as an underlying increase in BTN numbers. To further confirm these results, we electroporated the *Lzts sgRNAs* alongside the *Asic>GFP* reporter, another published marker of the BTNs^1,33^. While control larvae had fewer than 4 GFP+ BTNs in the majority of the larvae, we observed a net increase in the number of BTNs in the *CRISPR* condition. More specifically, nearly 50% of larvae had 4 or more GFP+ BTNs, again suggesting the generation of supernumerary BTNs (**Figure 4C,D**). This was further confirmed by endogenous *Asic* mRNA *in situ* hybridization which revealed the presence of supernumerary *Asic*-expressing cells expressing in the crispants (**Figure 4E, Supplementary Figure 9**). In some cases, supernumerary BTNs appeared to migrate ventrally or laterally instead of dorsally.

**Figure 4.**
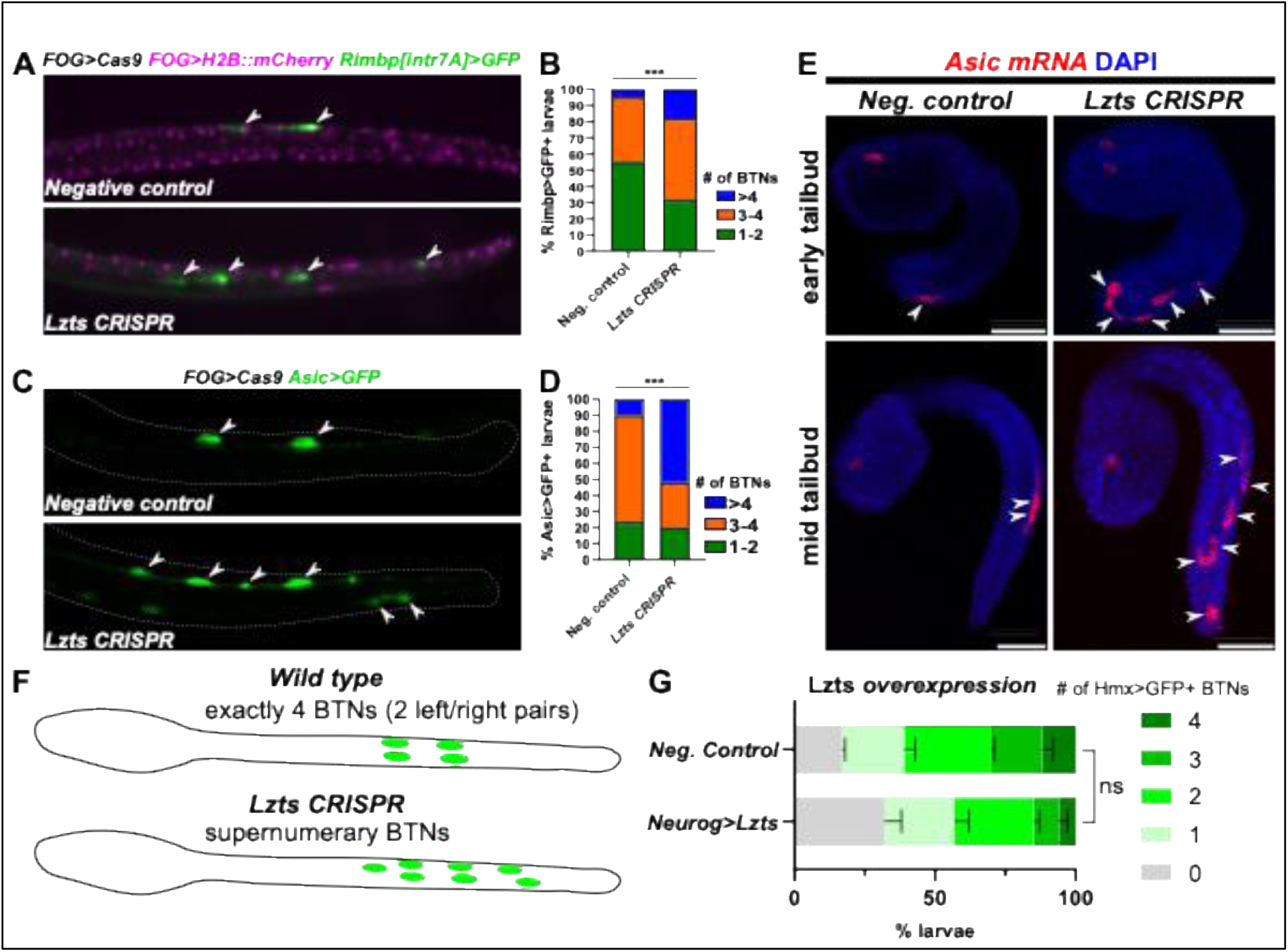
*Ciona Lzts* CRISPR knockout experiments. **A)** *Lzts* gene inactivation by CRISPR/Cas9 increases the normal number of BTN cells (labeled with *Rimbp[Intr7A]>GFP*, white arrows). FOG>H2B::mCherry labels cells where Cas9 is expressed (magenta nuclei). **B)** Graph showing a significant increase of BTN cells in the crispant larvae as seen in panel A (n = 161 & 153, respectively). Note that not all BTNs are visible in any given larva due to mosaic uptake of electroporated reporter plasmid. p****<0.0001. **C)** The same increase in BTNs is seen using a different plasmid reporter (*Asic>Unc-76::GFP*, in green). **D)** Graph showing a significant increase of BTN cells in the crispant larvae as seen in panel C (n = 110 & 101, respectively). p****<0.0001. **E)** *Asic* mRNA *in situ* hybridization (ISH, red) reveals the same increase in BTN numbers during development upon *Lzts* CRISPR (white arrows). Embryos counterstained with DAPI (blue). *FOG>Cas9* was used as before. Scale bars = 50 μm. **F)** Schematic diagram summarizing the *Lzts* CRISPR phenotype resulting in supernumerary BTNs instead of the invariant wild type condition of exactly 4 BTNs (two left/right pairs). Axons omitted for clarity. **G)** Scoring of BTN numbers comparing *Lzts* overexpression to negative control condition. The modest reduction in BTN numbers was not deemed statistically significant, by Fisher’s exact test, as no pair-wise comparisons between different BTN numbers were significant in both duplicates. n = 50 for each condition per duplicate.

We further replicated the *Lzts* CRISPR using the *GAD>GFP* reporter (**Supplementary Figure 10**), which is expressed in the GABAergic aBTNs, but not in the pBTNs^34,35^. Our results showed that in the CRISPR condition there was an increased number of larvae with two *GAD>GFP+* aBTNs. In contrast, in the control conditions we observed that the majority of larvae had only one aBTN labeled and ∼10% had two aBTNs labeled (**Supplementary Figure 10**), reflecting the typical mosaicism of plasmid uptake following electroporation. Thus, through multiple readouts we confirmed that the *Lzts* gene is required to control the number of BTNs, and that loss of Lzts function increases the number of BTNs, including aBTNs, generated in the *Ciona* larva (**Figure 4F**). Because *Lzts* loss-of-function results in supernumerary BTNs, we also asked whether overexpression of Lzts could suppress BTN numbers. When we overexpressed Lzts specifically in the BTN lineage using the BTN-specific *Neurog* enhancer (*Neurog[BTN]>Lzts*), there was a slight, but statistically insignificant reduction of cells labeled by yet another robust BTN marker, *Hmx>GFP* **(Figure 4G**). Taken together, these data suggest that Lzts is necessary to prevent supernumerary BTNs, but its overexpression may not be sufficient to reduce normal BTN numbers.

### *Lzts* CRISPR does not expand *Neurog* expression

Since the role of Neurog is crucial for the specification of BTNs^1,2^ and it is conserved in its vertebrate orthologs^36,37^, we aimed to evaluate the effects of *Lzts* CRISPR and overexpression on the expression pattern of *Neurog*, in order to understand the placement of *Lzts* within the GRN underlying the formation of BTNs. Since Neurog overexpression can result in supernumerary BTNs^1,2^, we asked if the increased BTN number in *Lzts* CRISPR larvae might depend on the expansion of the *Neurog+* neurogenic BTN cell lineage. By *in situ* hybridization (ISH) of *Neurog* around the time of BTN specification (early/mid tailbud), we found no substantial difference between the control and *Lzts1* knockout, at least at this developmental stage (**Figure 5A**). When we analyzed the expression of a *Neurog>mCherry* reporter plasmid at the mid-tailbud stage, we observed supernumerary BTNs dividing while delaminating and migrating, consistent with findings in mammals^8^. However, we did not detect general expansion of *Neurog* expression beyond the normal neurogenic territory of the tail tip (**Figure 5B**). Taken together these data suggest that *Lzts* is not required for precise expression of *Neurog* in the *Ciona* embryo, and that the supernumerary BTNs we observe in *Lzts* CRISPR larvae must be due to a different mechanism other than expanded *Neurog* expression.

**Figure 5.**
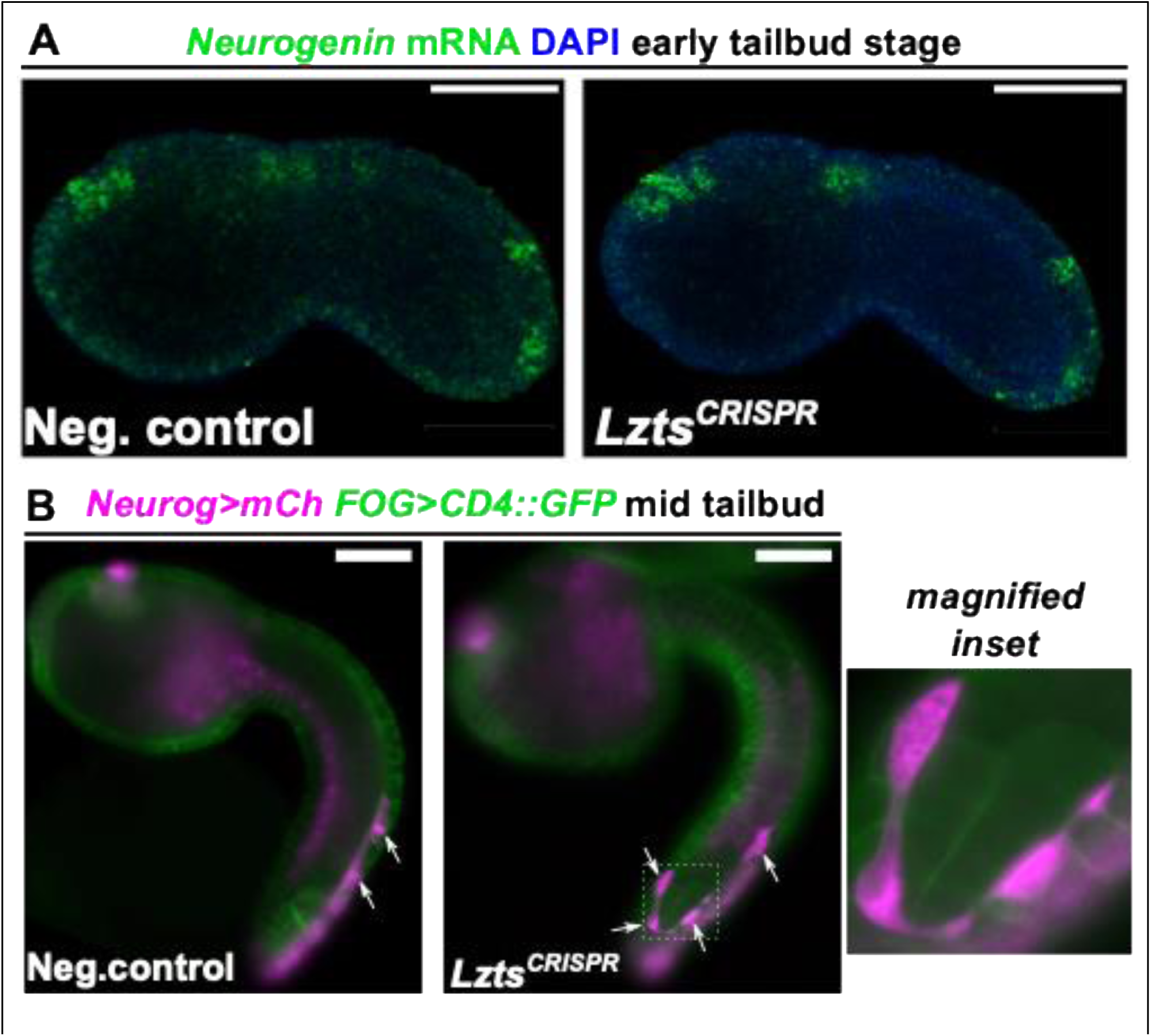
*Lzts* CRISPR does not expand the expression domain of the BTN regulator Neurogenin. **A)** mRNA *in situ* hybridization of *Neurog* does not reveal its expansion beyond the normal neurogenic cells of the tail tip in *Lzts* CRISPR early tailbud embryos when compared to negative control embryos. **B)** Assaying a *Neurog>Unc-76::mCherry* reporter slightly later also indicated no expansion of *Neurog* expression domain, but did reveal the same supernumerary migrating BTN precursors (white arrows) as seen with *Asic in situ* hybridization (see Figure 4E). As seen in the magnified inset, BTN precursors appear to still be dividing while migrating, and migrating ventrally instead of dorsally. Tissue-specific CRISPR was performed using *FOG>Cas9*.

### Proper BTN numbers can be restored in *Lzts* CRISPR larvae by concurrent manipulation of the Cdc25-Cdk1/2/3 pathway

Previous work in mammalian cell culture showed that Lzts1 accelerates mitosis through impaired Cdc25C phosphatase activity on Cdk1^5,16,38^. The current model proposes that Cdc25C-Cdk1 interaction requires Lzts1, and that absence of Lzts1 destabilizes Cdc25C. Thus loss of *Lzts1* results in an excessive increase of Cdk1 phosphorylation, lowering its activity and accelerating mitosis, through loss of M phase checkpoint activity and faster progression through prophase and prometaphase^5^. In mammalian *Lzts1-/-* cells, the normal, slower pace of mitosis was restored by co-expression of Cdc25C or a non-phosphorylatable form of Cdk1 (T>A/Y>F mutant). We therefore asked whether the same manipulations could restore or approximate normal BTN numbers in a *Ciona Lzts* CRISPR background. Indeed, overexpression of either *Ciona* Cdc25 or the predicted equivalent of non-phosphorylatable Cdk1 (*Ciona Cdk1/2/3^T>A/Y>F^)* brought BTN numbers down closer to that of negative control larvae (**Figure 6A-C**). This clear rescue effect was observed with either Asic>GFP- or Hmx>GFP-labelled BTNs. Our results suggest that Lzts regulates BTN numbers through a conserved cellular mechanism impinging on Cdc25/Cdk-mediated mitotic control (**Figure 6D**).

**Figure 6.**
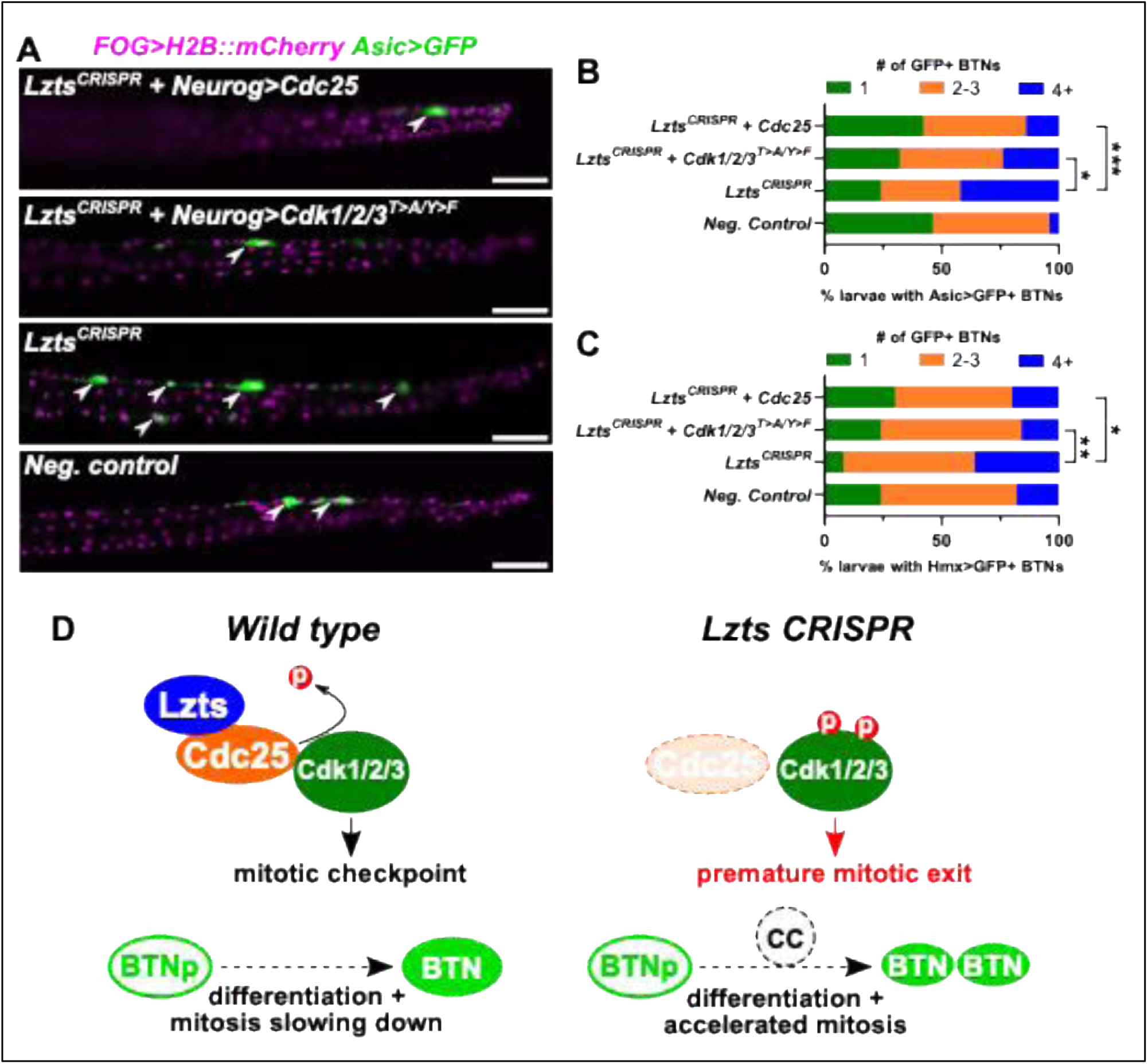
Overexpression of *Cdc25* or *Cdk1/2/3* phosphomutant rescues lower BTN numbers in *Lzts* CRISPR. **A)** Representative images of *Lzts* CRISPR larvae with or without overexpression of Cdc25 or a phosphomutant of Cdk1/2/3 (Cdk1/2/3^T>A/Y>F^) and compared to negative control larvae (U6>Control sgRNA used instead). *FOG* promoter was used to drive expression of Cas9 in the early BTN lineage, marked by FOG>H2B::mCherry (magenta). A BTN lineage-specific *Neurog* driver was used to overexpress Cdc25 or Cdk1/2/3^T>A/Y>F^. BTNs were visualized by *Asic>Unc-76::GFP (Asic>GFP)* reporter expression (green). Scale bars = 50 μm. **B)** Scoring larvae represented in panel A, binning into larvae with 1, 2-3, or 4+ Asic>GFP+ BTNs visible, showing that overexpression of Cdc25 or Cdk1/2/3 phosphomutant results in significant decrease of supernumerary BTNs in the *Lzts* CRISPR background. **C)** Rescue experiment replicated as in panels A and B, but using a different BTN reporter plasmid, *Hmx>Unc-76::GFP* instead. n = 50 for all conditions in panels B and C. * 0.03 < p < 0.05, ** p = 0.0116, *** p = 0.0017, all by Fisher’s exact test (one-tailed) comparing proportion of larvae with 4+ BTNs to the rest. GFP sequences tagged with N-terminal Unc-76. **D)** Proposed model and molecular pathway of Lzts function in BTN number control, based on mechanisms proposed by Vecchione et al. 2007. In wild type cells, Lzts normally stabilizes Cdc25 phosphatase, which dephosphorylates Cdk1/2/3 to ensure prolonged prophase and prometaphase as a mitotic checkpoint. In the absence of Lzts, Cdc25 is degraded, resulting in inactivation of Cdk1/2/3 and premature progression through prophase and accelerated mitosis. We propose that Lzts acts to slow down or arrest mitosis as BTN precursors differentiate, and that loss of Lzts results in a faster cell cycle (CC) and improper completion of mitosis during BTN differentiation, resulting in supernumerary BTNs.

## Discussion

Our results identify *Ciona* Lzts as a regulator of the bipolar tail neurons (BTNs) number and place this function within a broader, evolutionarily conserved picture of Lzts1/Fez1 biology. Loss of *Lzts* in the BTN lineage produced an excess of BTNs, likely through faster progression of precursor cells through mitosis. This accelerated cell division phenotype resembles the original description of *Lzts1-* null mouse embryonic fibroblasts^5,16,39^, in which loss of *Lzts1* destabilized Cdc25C, increased Cdk1 activity, and accelerated progression through prophase and prometaphase. In the mammalian fibroblast context, this acceleration came at expense of mitotic fidelity, causing chromosome missegregation and tumor predisposition^5,16,38^. In the *Ciona* BTN lineage, the most obvious consequence of accelerated mitosis appears to be an additional round of division that the lineage would not normally undergo, yielding supernumerary neurons (**Fig. 4F,G**). This contrasts with other perturbations, such as *Neurog* overexpression^1^, which specifies supernumerary BTNs by converting epidermal cells to a BTN fate. In our model (**Figure 6D**), a BTN is specified and starts to differentiate, but if mitosis proceeds more quickly than differentiation in the absence of Lzts, it would result in two BTNs per side instead of one.

Since the effect of *Lzts* CRISPR was mitigated by overexpression of Cdc25C or the Cdk1/2/3 phosphomutant, we suggest an ancestral core molecular function of Lzts as a “brake” on Cdk1/Cdc25C-dependent mitotic timing in the developing nervous system of tunicates. Our interpretation is supported by the observation that the main functional domains crucial for the interaction with Cdk1/Cdc25C (such as phosphorylation sites and leucine zipper motifs)^5,39^ are conserved between *Ciona* and mammalian Lzts proteins (**Supplementary Figure 11**). Furthermore, a conserved role for Lzts downstream of Neurog is consistent with previously published single -cell profiling^40,41^ from the developing mouse cortex (**Supplementary Figure 12A**), which shows high *Lzts1* expression levels in both Neurog2+ intermediate progenitors and migrating neurons for *Lzts1,* and negligible expression in still-cycling apical progenitors. Additionally, we found similar expression dynamics in published human single cell data^42^ **(Supplementary Figure 12B, Supplementary Figure 13B**). Interestingly, high coincident expression of *Lzts1* and *Neurog2* in intermediate progenitors is coherent with both our proposed model and previous findings on the role of vertebrate Neurog2 in driving cell cycle exit in neurons^43,44^. Then, we suggest the existence of a conserved mechanism regulating the cell cycle in neurons, with Neurog2 as a major regulator, in part through its ability to activate *Lzts1* expression.

Since the other three vertebrate orthologs (*Lzts2*, *Lzts3*, *Lzts4*) do not show significant expression levels in the aforementioned brain cell populations, we hypothesise that the original role of *Ciona Lzts* in the nervous system and cell cycle control in vertebrates has been retained primarily by *Lzts1*. The functions of the other *Lzts* paralogs in vertebrates may be related to the yet unknown functions of *Ciona Lzts* in other tissues (e.g. notochord). This hypothesis is supported by a major degree of conservation of functional domains with mammalian Lzts1 (**Supplementary Figure 11; Supplementary File 3**). In the invariantly developing *Ciona* embryo, Lzts may function as molecular “brake” on mitosis to ensure precise numbers of specific differentiated cell types, such as the BTNs. *Lzts* expression was also observed in other *Ciona* embryonic tissues, some of which also feature an invariant number of cells (e.g. the notochord, with its exactly 40 cells). However, the results of overexpressing Lzts were not quite as clear, since its overexpression did not substantially reduce BTN numbers. It is possible that increasing Lzts levels might not result in a proportional slowdown of mitosis, but rather the cell cycle is delayed by a simple presence or absence of *Lzts*.

Our findings reveal an *in vivo* developmental role for the Lzts-Cdc25-Cdk1 pathway that has not emerged from studies of mammalian corticogenesis. Although Lzts1 has been shown to stabilize Cdc25C and promote Cdk1 activity in cultured cells and mouse embryonic fibroblasts^5,16,38^, developmental studies in the mouse cortex have primarily highlighted its roles in cytoskeletal regulation, neuronal delamination, and progenitor positioning^7–9^.

Possibly, the loss of the cytoskeletal functions of Lzts1 dominates the mutant phenotype of mammalian cortex, and any effect on cycle length may be too subtle, or too distributed across a large, heterogeneous and regulative progenitor pool to register as a population-level change in proliferation. In the *Ciona* BTN lineage, even a modest change in the duration of mitosis is sufficient to add an extra, otherwise-absent division, and is therefore directly visible as a change in cell number. However, in multiple cases we noticed supernumerary BTNs situated ventrally instead of dorsally (**Figure 4C**), which appeared to be due to change in direction during migration (**Figure 4E, 5B**). We speculate that proper delamination and directed migration require mitotic arrest, and that delaminating/migrating BTNs undergoing an extra round of cell division can be misrouted as a result. This may connect the general function of Lzts proteins in cell division control and their specific effect on neurodevelopment.

More broadly, our findings support the idea that the molecular logic linking cell-cycle regulators to neuron number might be like the logic that links those regulators to tumor suppression and tissue architecture (**Figure 6**). We propose that the Lzts/Cdc25c/Cdk1 axis may be one of the most deeply conserved regulatory nodes in bilaterian cell-cycle control. In the context of previous work on BTN specification, differentiation, we can link the BTN gene regulatory network to a conserved mitotic-timing factor for precise neuron number control. Future work should address whether *Ciona* Lzts also regulates cell number in its other expression domains, i.e. the notochord and stomodeum. Furthermore, additional approaches might open up new avenues in understanding the novel role of *Lzts* ohnologs in corresponding cells/tissues of vertebrates. It also remains to be seen if *Ciona* Lzts influences the cytoskeletal and spindle positioning behaviors documented for its mammalian counterpart in the developing cortex. Such experiments would help establish whether the cytoskeletal-remodeling functions of Lzts proteins were present in the last common ancestor of tunicates and vertebrates, or if novel functions arose as a consequence of subfunctionalization of different *Lzts* paralogues in jawed vertebrates. Additionally, they may help to reveal whether the proliferation and cytoskeletal functions of Lzts1 are mechanistically coupled through the same pathways, or if these involve different interaction partners in different cellular and/or subcellular contexts.

## Ethics statement

Ethical approval was not required because only invertebrate animals were used in this study.

## Acknowledgements

We thank Dr Kwantae Kim and Dr. Katarzyna Piekarz for technical assistance. Work in the Ristoratore lab was funded by the Assemble grant 730984. Paola Olivo was funded by a SZN-OU Phd fellowship and from a Travel Fellowship from “The Company of Biologists, Development” (DEVTF2110636). Work in the Stolfi lab was funded by grant R01HD104825 from NICHD. Work in the Coppola lab was funded by Florida Gulf Coast University (FGCU) startup funds (PI: Ugo Coppola).

## Materials and Methods

### Genome database searches and phylogenetic reconstruction

*Ciona robusta* Lzts protein sequence was used as a query in BLASTp and tBLASTn in genome databases of selected species (NCBI, Ensembl, Ensembl Metazoa, ANISEED^45^. The entire dataset of protein sequences (**Supplementary File 1**) for domain architecture was analyzed by using the domain database provided by Ensemb, and manually annotated. All the surveyed sequences were verified to be Lzts proteins through BLASTs and domain analyses. The surveys were calibrated with multiple species from agnathans to primates to take into account the impact of multiple WGDs in vertebrates^18,19^ and in teleosts^23,46^. Orthology of the Lzts sequences was initially assessed by using a reciprocal best blast hit (RBBH) approach employing default parameters and corroborated by phylogenetic analyses. Protein alignments for phylogeny were generated employing Clustal Omega^47^. The phylogenetic reconstruction of Figure 1 was carried out on the total protein sequences and based on maximum-likelihood (ML) inferences calculated with PhyML3.0^48^, employing automatic Akaike Information Criterion (AIC) by Smart Model Substitution (SMS)^49^, which selected the JTT+G+F model employing discrete gamma distribution in categories. All parameters (gamma shape = 0.8; proportion of invariants (fixed) = 0.000) were calculated from the dataset. Branch support was provided by aLRT^50,51^. The phylogeny of S1 Fig was carried out employing Bayesian Information Criterion (BIC) by SMS, which sorted the JTT+G+F model using discrete gamma distribution in categories. All parameters (gamma shape = 0.8; proportion of invariants (fixed) = 0.000) were established from the dataset, with aBayes branch support. The alignment of selected Lzts proteins in Supplementary Figure 10 was performed employing Clustal Omega^47^ with the domains that were manually mapped. The presence of domains was assessed using Ensembl and PROSITE^52^ datasets, plus confirmed with previous analyses^16^.

### Intron/exon structure analysis

Gene structures were classified merging the genomic sequences with ESTs when available, as previously described^26,53–55^. Introns were classified as phase 0, phase 1, and phase 2, according to their positions within the protein-reading frames. The amino-acids containing the conserved introns represented in Figure 1B were manually mapped on a Clustal Omega alignment^47^ of selected Lzts proteins (**Supplementary File 2**).

### Syntenic analysis

The presence/absence of synteny was assessed by examining the chromosomes on public genome databases (NCBI, Ensembl, Ensembl Metazoa, ANISEED), and confirmed using the Genomicus database^56^. The window considered for the loci analysis consisted of twenty flanking genes. Not conserved genes were not included from the analysis. All the genes were represented employing colored rectangles, using the same color for all *Lzts* genes (red).

### Regulatory analysis and molecular cloning

The *Ciona robusta* regulatory region of Figure 3 was aligned to ATAC-seq peaks^28,57^. Genomic DNA sequences were retrieved from Aniseed^45^, GHOST^58^, and Ensembl databases. The *cis*-regulatory elements upstream *Lzts* were PCR-amplified from genomic DNA the products were inserted into GFP^59^ or Unc-76::GFP containing vectors^60^, with or without a basal promoter of *FOG* ^30^, or custom-synthesized and cloned by Twist Bioscience. *Lzts, Cdc25,* and Cdk1/2/3 (phosphomutant) coding sequences were custom-synthesized and cloned by Twist Bioscience. All relevant recombinant DNA sequences can be found in **Supplemental File 4.**

### Electroporations

Adults of *Ciona robusta* were collected from the Gulf of Naples, or from San Diego, CA, United States, by M-REP or Marinus. The gametes from multiple animals were gathered separately for *in vitro* cross-fertilization, subsequently followed by dechorionation and electroporation as previously illustrated^28,29^. Electroporated plasmids (e.g., 10 μg) were 700 μl of total volume. Embryos were staged following the developmental timeline ^61^. In order to visualize fluorescent reporters (GFP, mCherry), embryos were fixed employing MEM-FA (3.7% methanol-free formaldehyde, 0.1 M MOPS pH 7.4, 0.5 M NaCl, 2 mM MgSO4, 1 mM EGTA) for 30 min and washed several times in PBS-NH4Cl and in PBS containing 0.05% Triton X-100. Each electroporation for CRISPR/Cas9 experiments was carried out using 60 μg of plasmid. The statistical significance of electroporations and perturbations were evaluated using Fisher’s exact test or g Chi-square test for trend. Fisher’s exact tests were calculated using GraphPad Prism’s online 2×2 contingency table analyzer: https://www.graphpad.com/quickcalcs/contingency1.

### *In situ* hybridization

Whole-mount fluorescent mRNA *in situ* hybridization experiments were performed out as described previously^62^, using DIG- and FLUO-labeled riboprobes combined with anti-DIG-POD or anti-FLUO-POD Fab fragments (Roche, Indianapolis, IN), and Tyramide Amplification Signal with Fluorescein (Perkin Elmer, MA). The antisense riboprobes were obtained from a PCR-amplified fragment, subsequently cloned in the TOPO-TA vector (Invitrogen). The template sequences are listed in **Supplementary File 4**.

### CRISPR/Cas9 approach

The selection and cloning of the sgRNA to use for CRISPR/Cas9-mediated mutagenesis has been performed following the protocol adapted from Gandhi et al., 2018. The predictive algorithm used for the design of the *in vivo*-transcribed *sgRNAs* has been the Fusi/Doench^63^, available on the CRISPOR portal^64^. The selected target sequences (**Supplementary File 4**) have been selected not too close to the translational start nor too far towards the C terminus, to have the higher impact on the function of protein of interest^31^ Then, the *sgRNA* plasmids were validated by electroporation following the aforementioned protocol, using 25–50 μg *U6>sgRNA* and 25 μg *Ef1α >nls::Cas9::nls* (per 700 μl electroporation volume). Collected larvae (50-100) were used to extract genomic DNA using the QIAamp DNA Micro Kit (Qiagen), followed by a PCR amplification of the target sequence. Genomic primers were designed to amplify a fragment 300–1500 bp long (Pfx Platinum, Thermo Fisher Scientific), with the target sites at least 150 bp away from each end of the fragment. The PCR products were purified using NucleoSpin Gel and PCR clean-up (Macherey Nagel), and analyzed for Sanger sequencing. The primers used for sequencing are the same used for performing the CRISPR mutagenesis.The efficiency of the *sgRNAs* was evaluated using the Tide algorithm^65^. For tissue-specific CRISPR knockout in the BTN lineage, the *FOG* promoter was used to drive expression of Cas9 or optimized Cas9::GemN^66^. For CRISPR/rescue experiments, 30 μg of *FOG>Cas9,* 40 μg of each sgRNA, 10 μg of *FOG>H2B::mCherry,* 70 μg of *Asic* or *Hmx>Unc-76::GFP,* and 70 μg of *Neurog[BTN]>Cdc25* or *Cdk1/2/3^T>A/Y>F^* were used per 700 μl total electroporation volume. For overexpression, 100 μg of *Neurog[BTN]>Lzts* and 35 μg of *Neurog[BTN]>H2B::mCherry* were used instead.

### Single-cell datasets analysis

Single-cell data from the mouse nervous system were retrieved from previously published datasets ^40,41^, using default functions (“NewCellType”) from the [Single Cell Portal](https://singlecell.broadinstitute.org/single_cell). Single-cell data from the human nervous system were retrieved from [SnMultiome](https://cell.ucsf.edu/snMultiome/), a standardized dataset^42^ of intermediate progenitors and migrating neurons.

