## supplementary figures for "Fine-tuning neuron number: *Lzts* as a key regulator of neurogenesis and cell proliferation in *Ciona*"

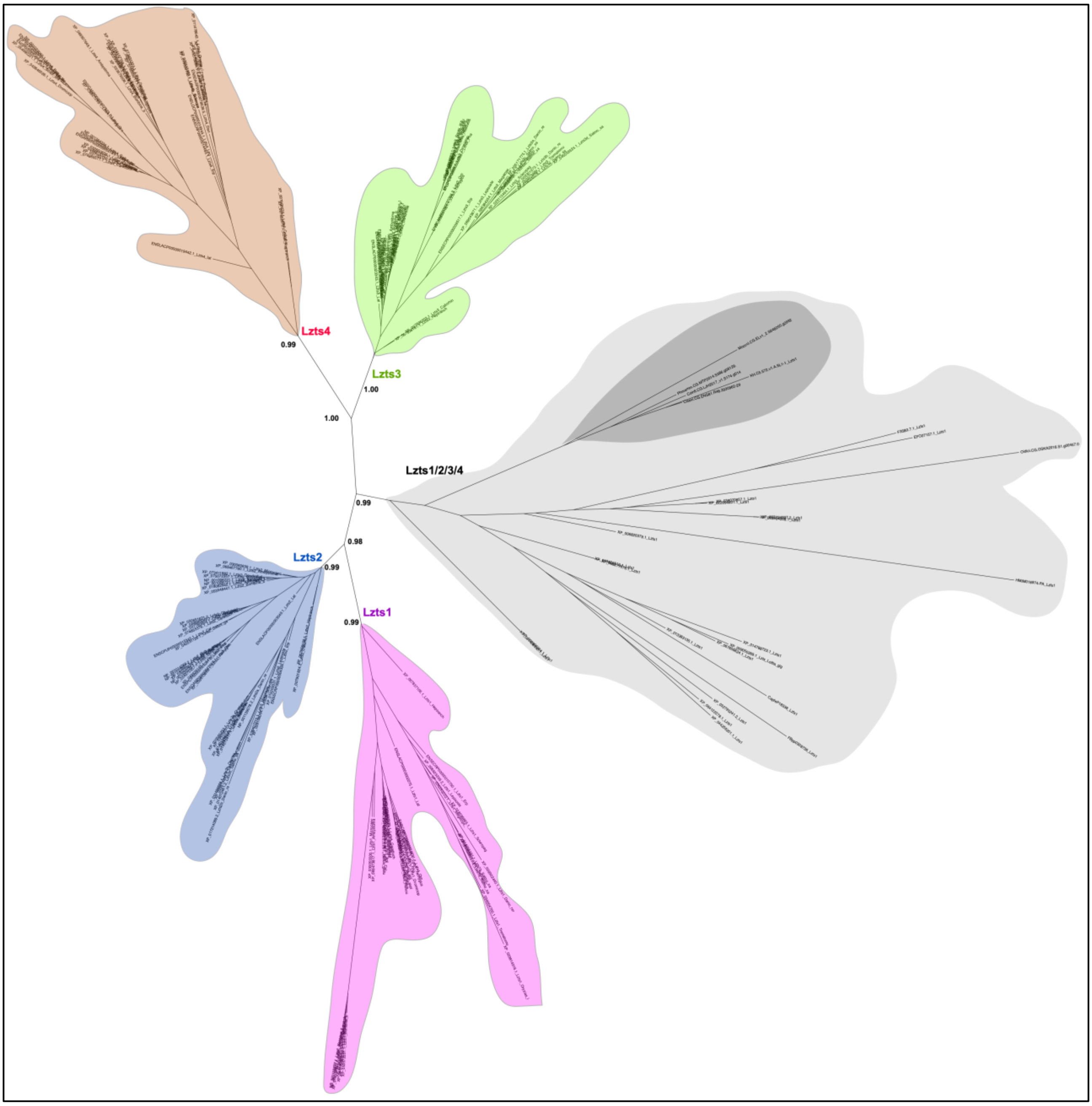


**Supplementary Figure 1.** Phylogenetic tree of Figure 1A (including the names of species and proteins), with numbers at the branches indicating replicates generated using aLRT method. The colors correspond to those used in Figure 1A.

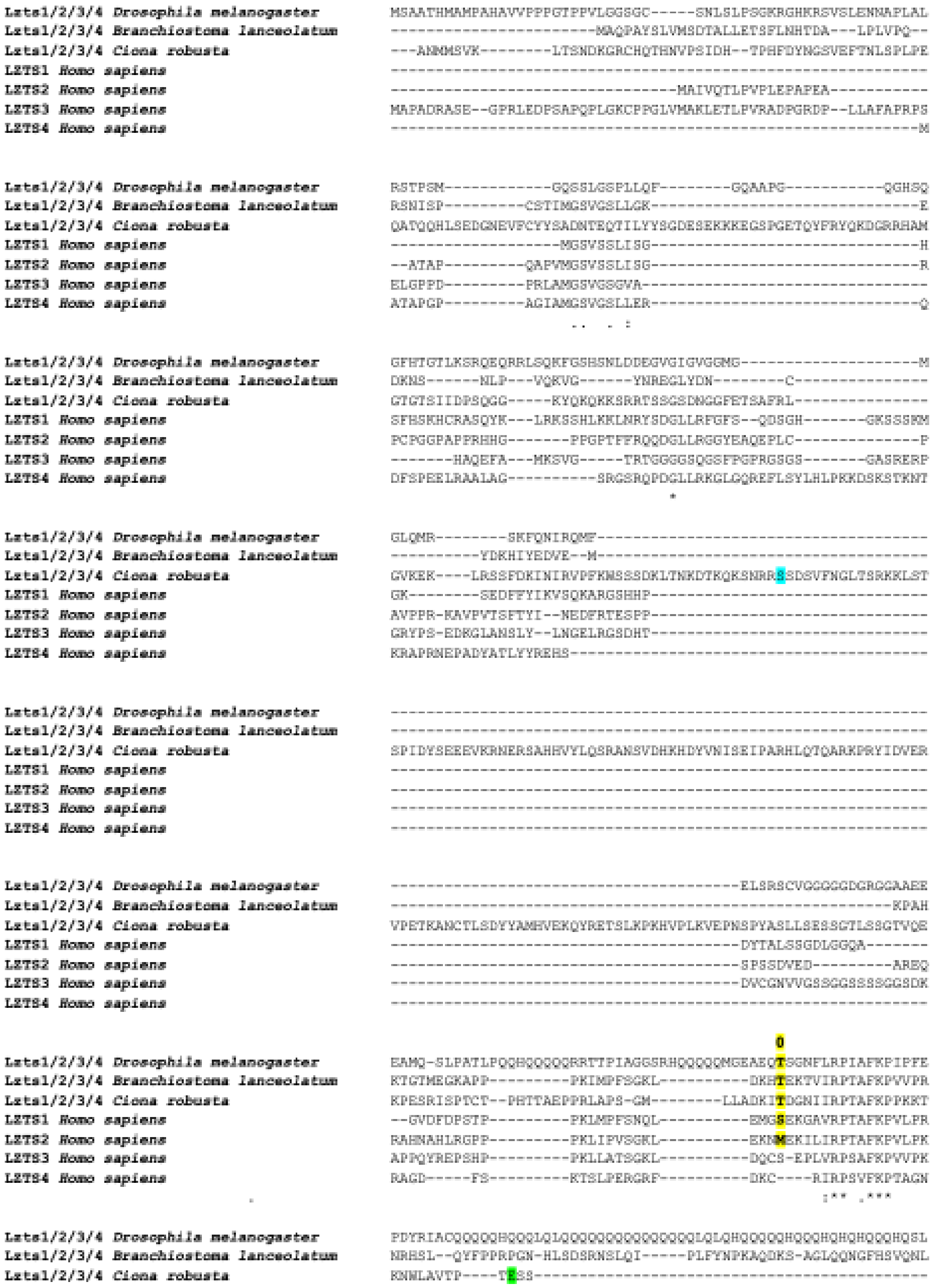


**Supplementary Figure 2.** Intron/exon junctions represented on a ClustalX alignment. The conserved junctions are highlighted using both bold and yellow (representing phase 0).

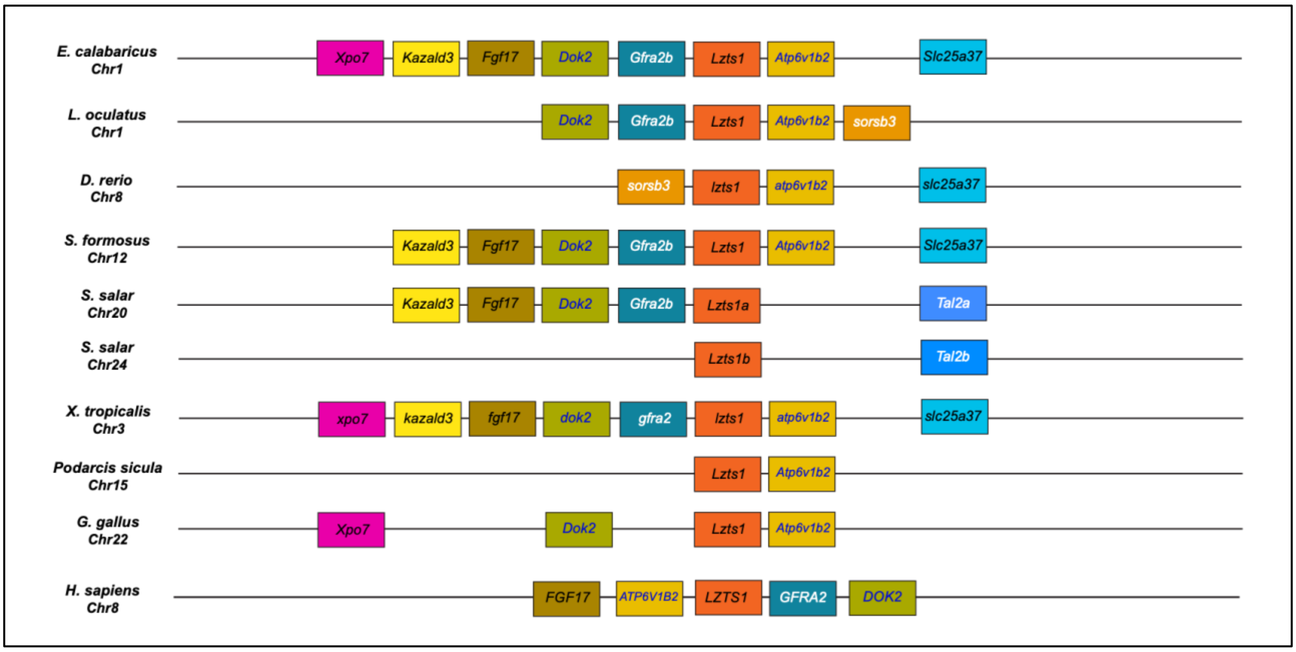


**Supplementary Figure 3.** Synteny analysis of vertebrate *Lzts1* gene locus. Schematization of conserved genomic environments of *Lzts1* (red rectangles) in selected vertebrate species with relative chromosomes/scaffolds information. Flanking orthologous genes are represented using rectangles of the same color

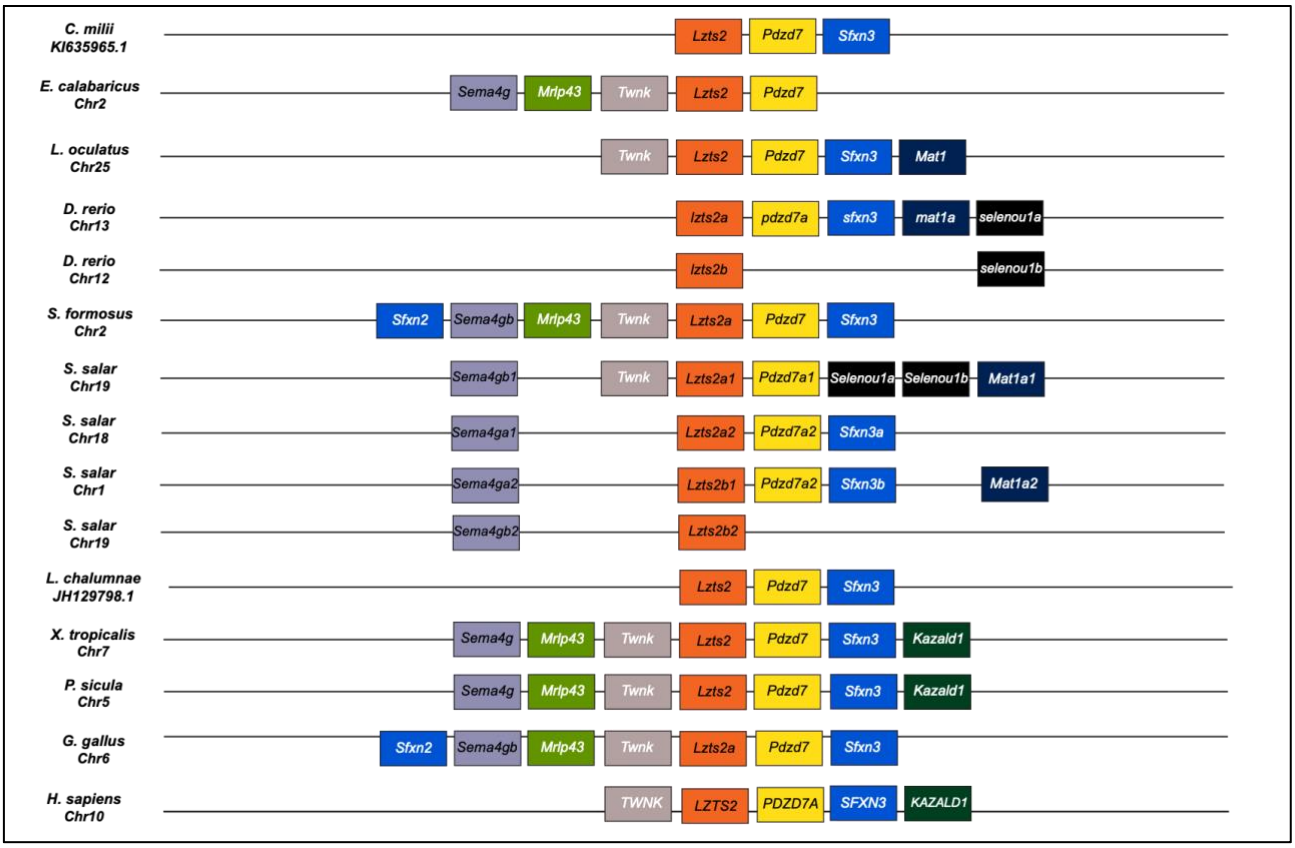


**Supplementary Figure 4.** Synteny analysis of vertebrate *Lzts2* gene locus. Schematization of conserved genomic environments of *Lzts2* (red rectangles) in selected vertebrate species with relative chromosomes/scaffolds information. Flanking orthologous genes are represented using rectangles of the same color.

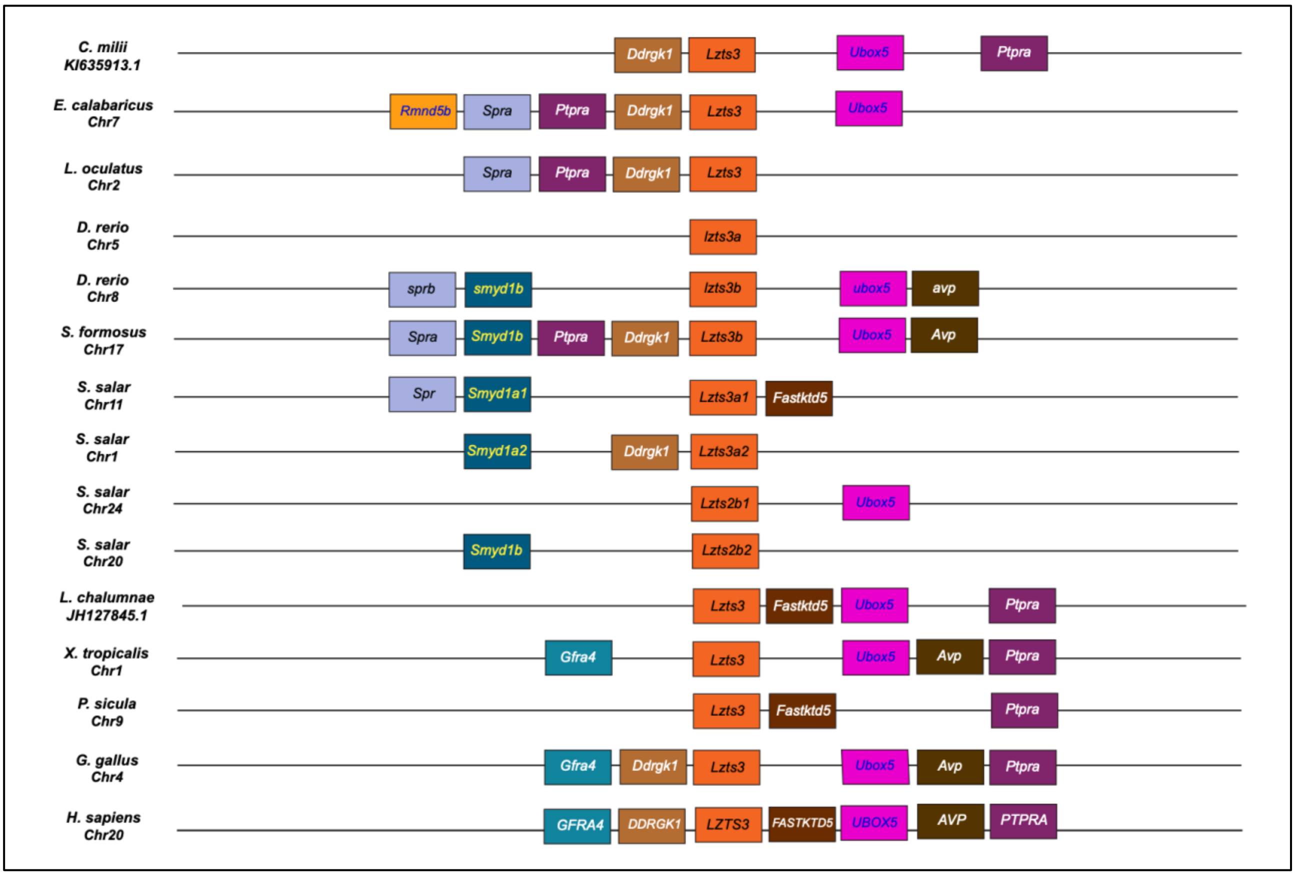


**Supplementary Figure 5.** Synteny analysis of vertebrate *Lzts3* gene locus. Schematization of conserved genomic environments of *Lzts3* (red rectangles) in selected vertebrate species with relative chromosomes/scaffolds information. Flanking orthologous genes are represented using rectangles of the same color.

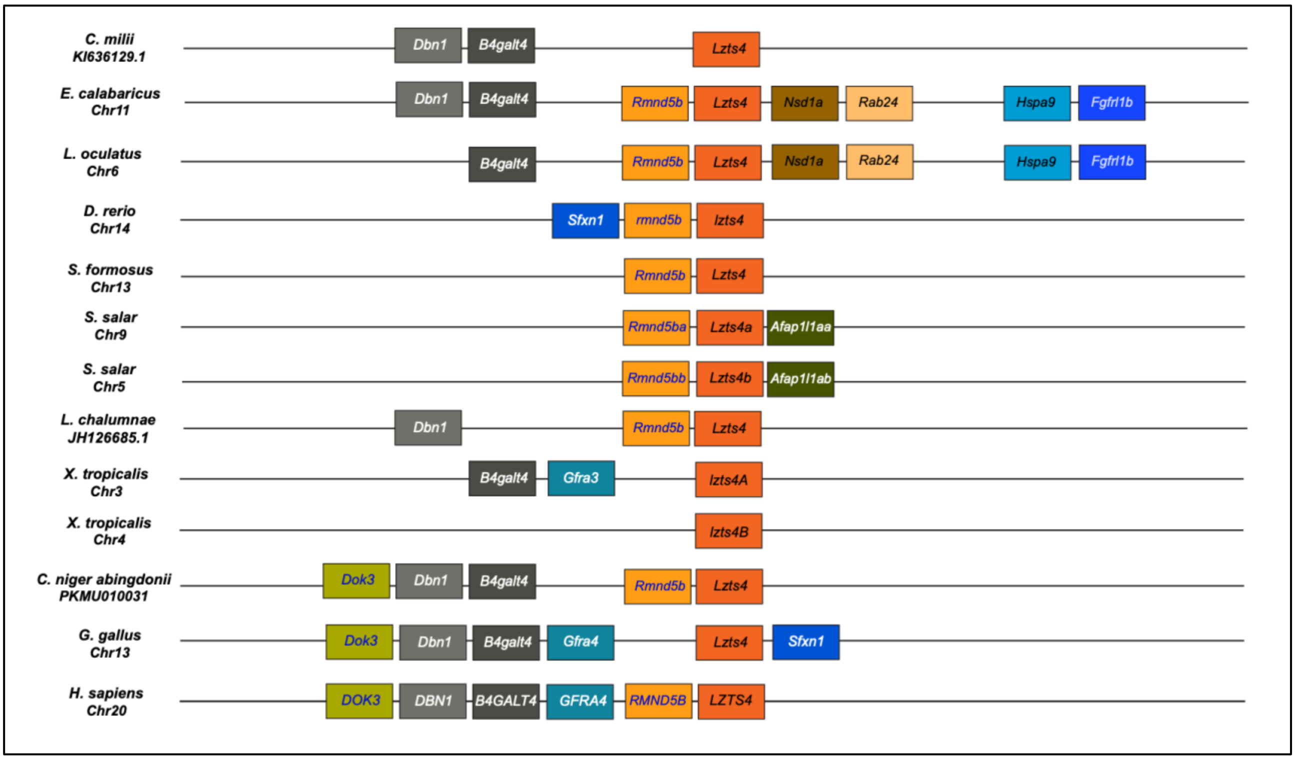


**Supplementary Figure 6.** Synteny analysis of vertebrate *Lzts4* gene locus. Schematization of conserved genomic environments of *Lzts4* (red rectangles) in selected vertebrate species with relative chromosomes/scaffolds information. Flanking orthologous genes are represented using rectangles of the same color

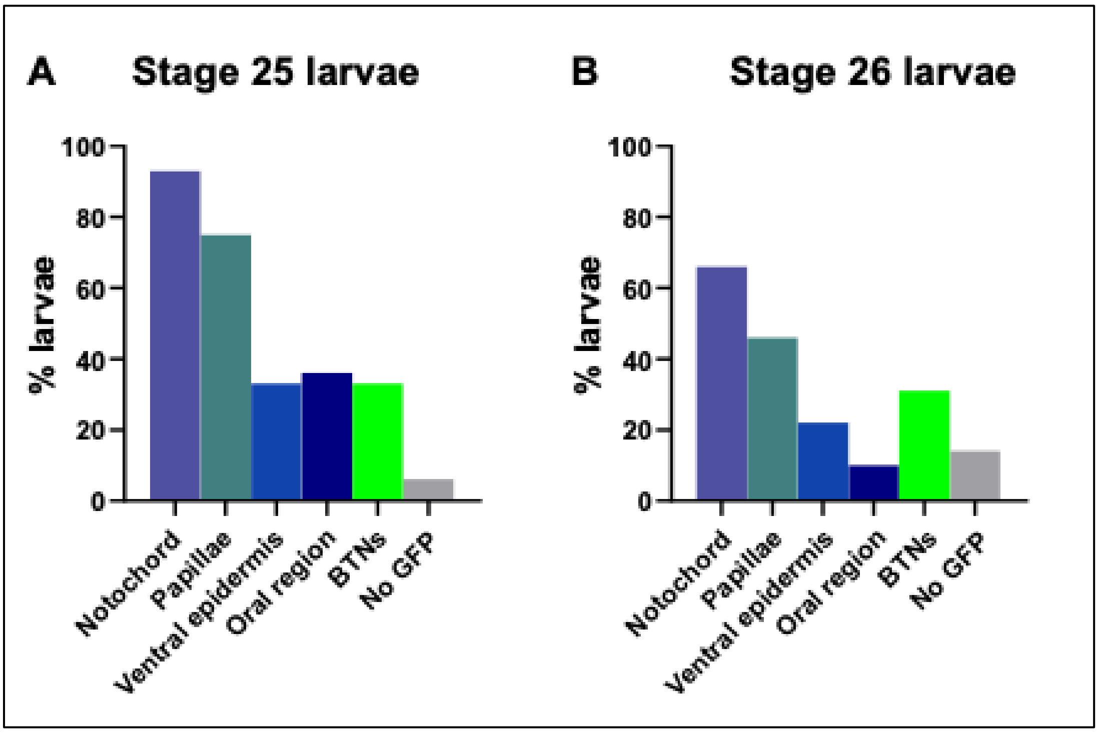


**Supplementary Figure 7. A, B)** Graphs showing the expression distribution of the *LztsA>GFP* reporter in larvae, at stages 25 and 26, respectively

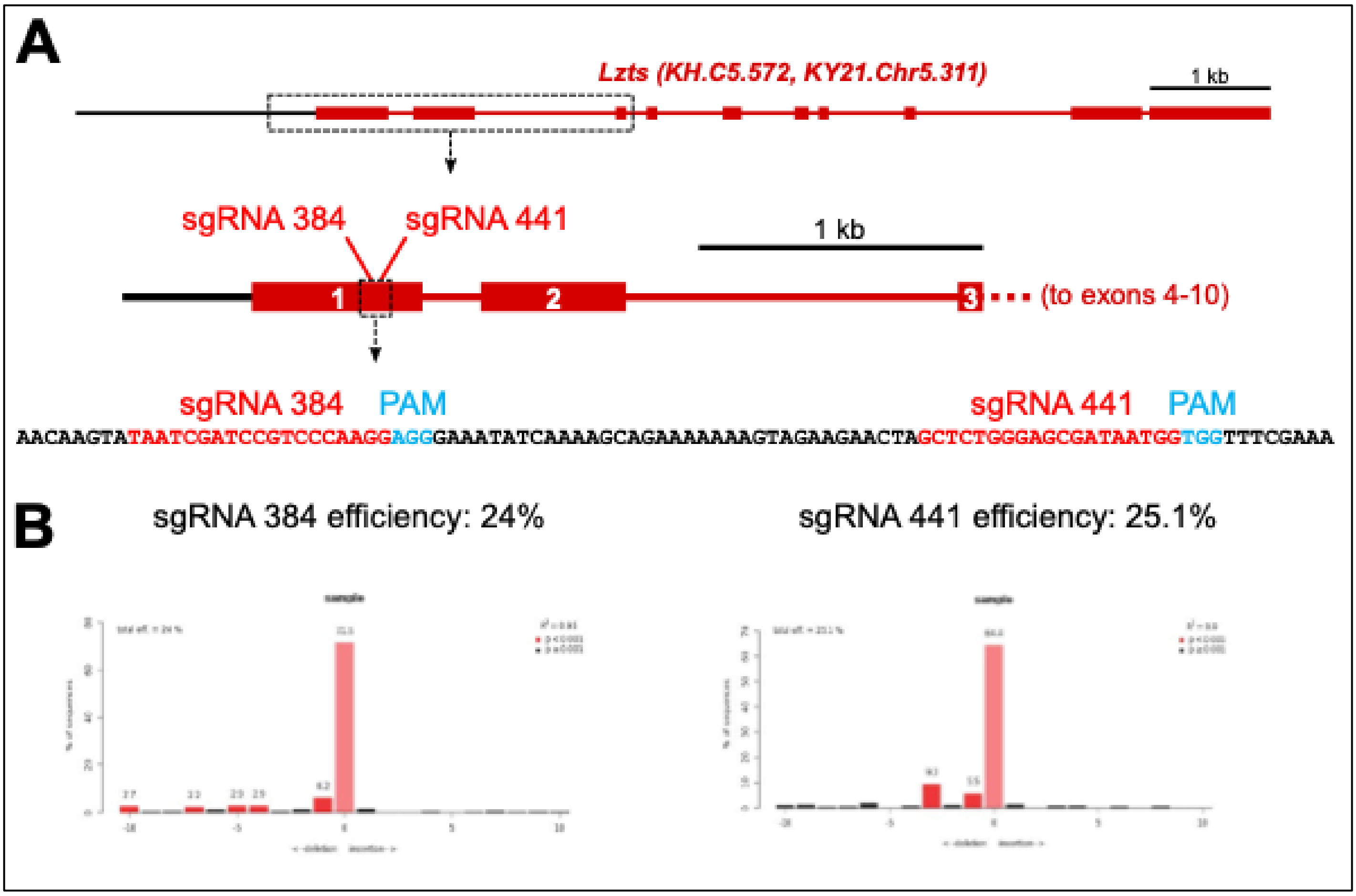


**Supplementary Figure 8.** Scheme showing the relative positions of sgRNAs used to perform the CRISPR/Cas9-mediated gene inactivation and its efficiency.

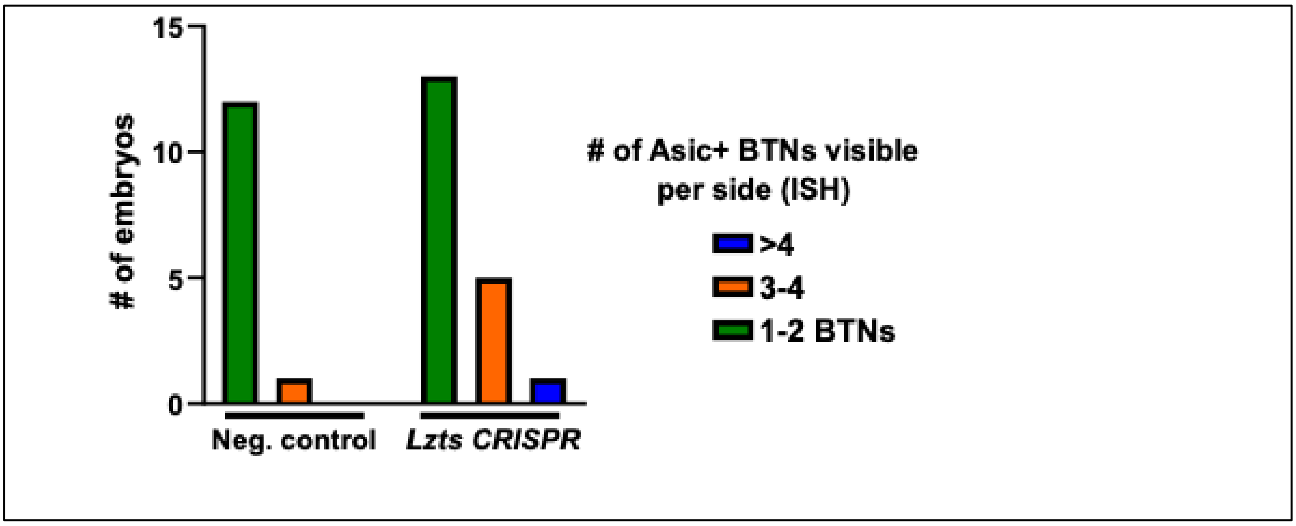


**Supplementary Figure 9.** Plot showing number of BTNs in *Lzts* CRISPR and negative control tailbud embryos, as assayed by *Asic in situ* hybridization, shown in Figure 4E. N = 24 (control) and 27 (CRISPR).

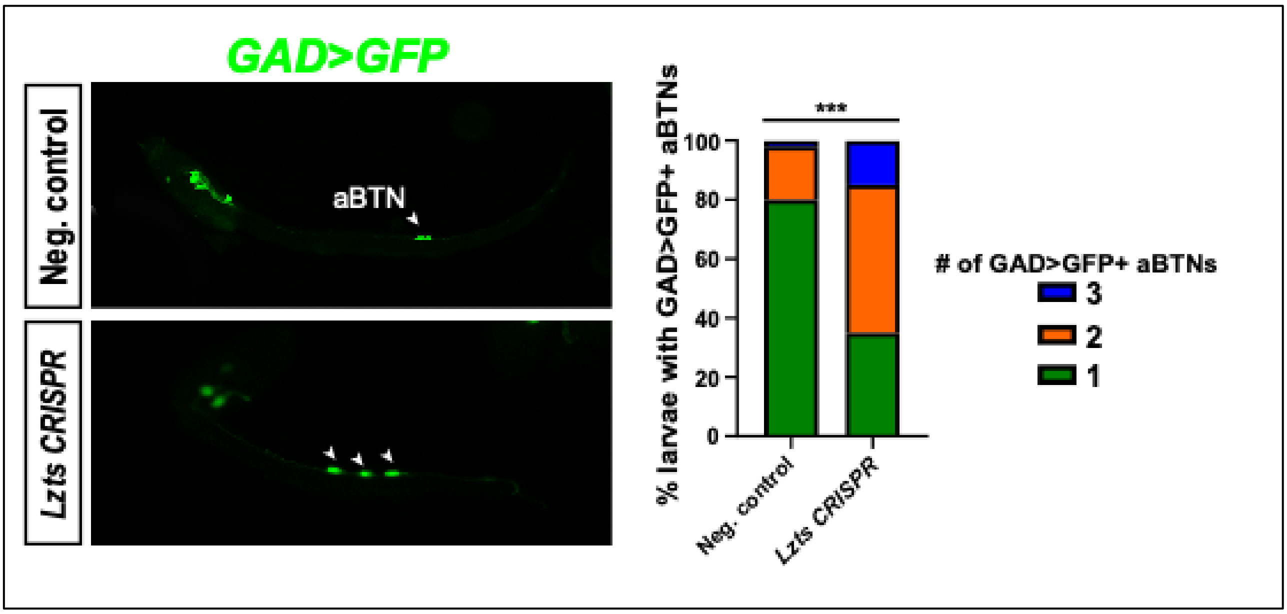


**Supplementary Figure 10.** Left: comparison between negative control and *Lzts* CRISPR larvae, assaying number of anterior BTNs (aBTNs) using *GAD>Unc-76::GFP* as marker. Right: graph indicating an increase in aBTNs in the crispant larvae. N = 108 (control) and 80 (CRISPR). Tissue-specific CRISPR was performed using *FOG>Cas9.* Fisher's exact test has been used on pairwise comparisons of all perturbations of aBTN number. *** $p < 0.0001$.

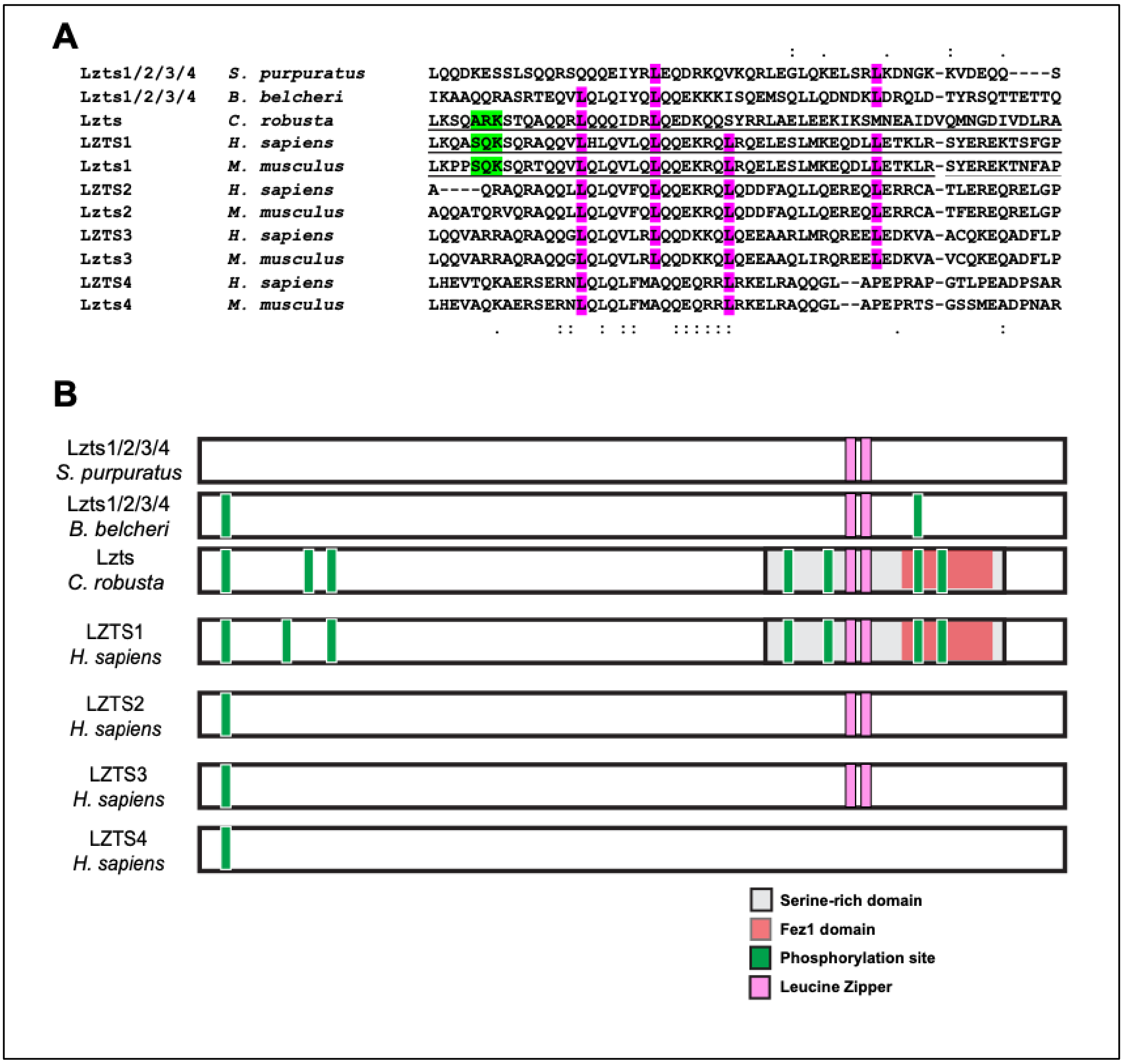


**Supplementary Figure 11. A)** Partial protein alignment showing conservation between *C. robusta* and other models of a phosphorylation site (green) and leucine zippers (magenta). **B)** Schematic with selected Lzts proteins including typical Lzts protein domains (in red Fez1 domain, grey the Serine-rich domain, in magenta the Leucine “zippers”) and in green the conserved hypothetical phosphorylation sites (adapted from[16]). Only the domains/sites detected in *C. robusta* were represented.

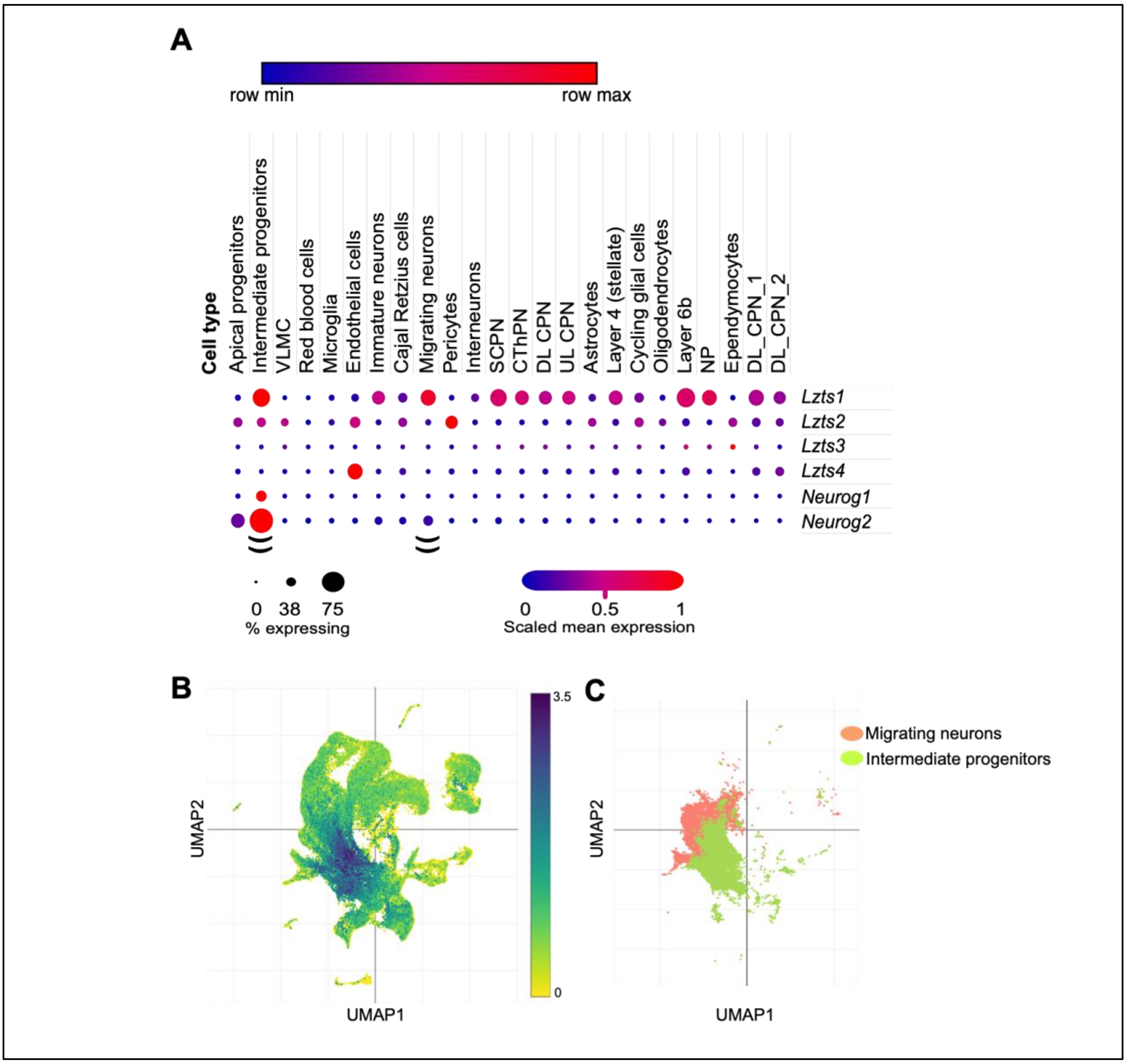


**Supplementary Figure 12. A)** Dotplot showing higher levels of expression of *Lzts1* and *Neurog2* in intermediate progenitors and migrating neurons (double parenthesis), from developing mouse cortex single-cell RNA sequencing data (see text for details). **B)** UMAP representation of relative co-expression of *Lzts1* and *Neurog2* from the same dataset. **C)** UMAP representation with the coordinates of the murine brain, highlighting intermediate progenitors and migrating neurons, which overlap with the greatest extent of *Lzts1* and *Neurog2* overlap in panel B. Plots generated using Single Cell Portal[40,41].

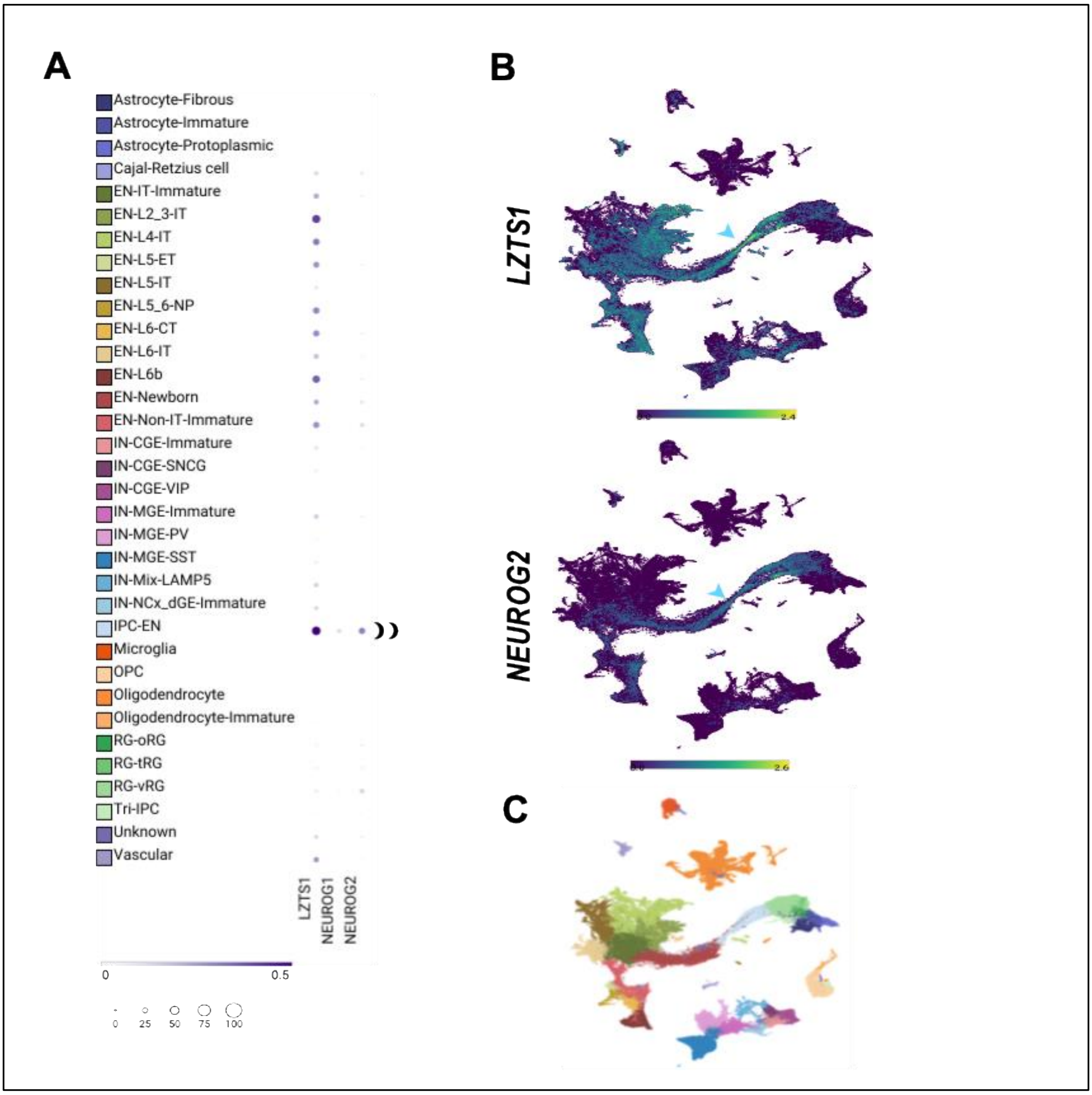


**Supplementary Figure 13. A)** Dotplot showing higher levels of expression of *LZTS1* and *NEUROG2* in intermediate progenitors (IPC-EN) from human first-second trimester datasets from single-cell RNA sequencing (double parenthesis). Data retrieved from[67]**. B)** UMAP representation showing the relative co-expression of *LZTS1* and *NEUROG2* in IPCs (intermediate progenitors, light blue arrowheads). **C)** UMAP legend with all the brain subtypes. Plots generated using SnMultiome[42].

**Supplementary File 1.** List of protein sequences used for Lzts phylogenetic analyses.
**Supplementary File 2.** Raw data representing intron code generation.
**Supplementary File 3.** Clustal alignment with selected Lzts proteins including typical Lzts protein domains (in red Fez1 domain, underlined the Serine-rich domain, in magenta the Leucine "zippers"). In green the conserved hypothetical phosphorylation targets (adapted from Vecchione et al., 2007).
**Supplementary File 4.** Supplementary sequence file containing all regulatory DNA, cDNA, sgRNA, and validation primer sequences used in this study.
